# CCNE1 amplification is synthetic-lethal with PKMYT1 kinase inhibition

**DOI:** 10.1101/2021.04.08.438361

**Authors:** David Gallo, Jordan T.F. Young, Jimmy Fourtounis, Giovanni Martino, Alejandro Álvarez-Quilón, Cynthia Bernier, Nicole M. Duffy, Robert Papp, Anne Roulston, Rino Stocco, Janek Szychowski, Artur Veloso, Hunain Alam, Prasamit S. Baruah, Alexanne Bonneau Fortin, Julian Bowlan, Natasha Chaudhary, Jessica Desjardins, Evelyne Dietrich, Sara Fournier, Chloe Fugère-Desjardins, Theo Goullet de Rugy, Marie-Eve Leclaire, Bingcan Liu, Henrique Melo, Olivier Nicolas, Akul Singhania, Rachel K. Szilard, Ján Tkáč, Shou Yun Yin, Stephen J. Morris, Michael Zinda, C. Gary Marshall, Daniel Durocher

## Abstract

Amplification of the gene encoding cyclin E (*CCNE1*) is an oncogenic driver in several malignancies and is associated with chemoresistance and poor prognosis. To uncover therapeutic targets for *CCNE1*-amplified tumors, we undertook genome-scale CRISPR/Cas9-based synthetic lethality screens in cellular models of *CCNE1* amplification. Here, we report that increasing *CCNE1* dosage engenders a vulnerability to the inhibition of the PKMYT1 kinase, a negative regulator of CDK1. To inhibit PKMYT1, we developed RP-6306, an orally bioavailable and selective inhibitor that shows single-agent activity and durable tumor regressions when combined with gemcitabine in models of *CCNE1*-amplification. RP-6306 treatment causes unscheduled activation of CDK1 selectively in *CCNE1* overexpressing-cells, promoting early mitosis in cells undergoing DNA synthesis. *CCNE1* overexpression disrupts CDK1 homeostasis at least in part through an early activation of the FOXM1/MYBL2/MuvB-dependent mitotic transcriptional program. We conclude that PKMYT1 inhibition is a promising therapeutic strategy for *CCNE1*-amplified cancers.

## Main Text

Amplification of the *CCNE1* locus on Chr 19q12 is prevalent in multiple tumor types, particularly in high-grade serous ovarian cancer (HGSOC), uterine tumors and gastro-oesophageal cancers where high cyclin E levels are associated with genome instability, whole-genome doubling and resistance to cytotoxic and targeted therapies (*1–4*). In HGSOC, *CCNE1* amplification is detected in ∼20% of tumors, in a manner largely mutually exclusive with homologous recombination deficiency and is enriched in platinum-refractory tumors (*2, 5*). The paucity of therapeutic options for *CCNE1*-amplified tumors makes the development of novel therapeutics that target this amplification a critical unmet need (*6*). Cyclin E itself is not considered to be a druggable target but its cognate cyclin-dependent kinase (CDK) CDK2 is. CDK2 inhibition shows promising activity in *CCNE1*-amplified cell lines (*7*) but CDK2 inhibitors with minimal cross-reactivity towards other members of the CDK family have yet to progress into clinical development. Since CDK2 phosphorylation regulates cyclin E levels via FBXW7-driven ubiquitylation (*8–11*), transient CDK2 inhibition could also lead to paradoxical stabilization of cyclin E. Thus, we surmised that a synthetic lethality approach (*12*) exploiting vulnerabilities caused by the increase in cyclin E levels, may provide much needed novel therapeutic options for *CCNE1*-amplified tumors.

To identify genetic vulnerabilities to the increased dosage of *CCNE1*, we developed an isogenic pair of cell lines that stably overexpress cyclin E from a *CCNE1-2A-GFP* fusion integrated into the genome of RPE1-hTERT *p53^-/-^* cells using piggyBAC transgenesis (Fig S1a). These cells are hereafter referred to as “*CCNE1-*high” and also stably expressed Cas9 (*13*). We selected and characterized two clones, C2 and C21, which showed phenotypes consistent with cyclin E overexpression such as accumulation of cells in early S phase, elevated DNA replication stress as measured by CHK1 S345 phosphorylation and MCM helicase loading defects (Fig S1a-c). We undertook genome-scale CRISPR/Cas9 screens in the parental and both *CCNE1*-high clones using the genome-scale TKOv2 single-guide (sg) RNA library (*14*) and also subsequently rescreened clone C2 with the TKOv3 sgRNA library, which has improved performance (*15*) (Fig. 1a). Using two CRISPR screen scoring methods, CCA (*16*) and BAGEL2 (*17*), we identified 5 genes whose mutation caused a selective loss of fitness in *CCNE1*-high cells, in all three screens: *ANAPC15*, *FBXW7*, *PKMYT1*, *UBE2C* and *UBE2S* (Fig. 1b, Table S1 and Datasets S1-S2). To prioritize this list, we mined the genetic dependencies of cancer cell lines derived from the DepMap project (*18*). This analysis identified *PKMYT1* as the only gene, among the 5 hits, which displayed a strong and selective dependency in *CCNE1*-amplified tumor cell lines (Fig 1c and Table S2). *PKMYT1* encodes an evolutionarily conserved protein kinase, also known as Myt1, whose primary role is the negative regulation of CDK1 both by its inhibitory phosphorylation on Thr14 and its sequestration in the cytoplasm (*19–23*). PKMYT1 is structurally related to – and much less studied than – WEE1, which phosphorylates the adjacent Tyr15 residue on CDK1 and CDK2 to inhibit these kinases (*24–26*). Unlike WEE1, which is nuclear-localized, PKMYT1 is sequestered in the cytoplasm by an interaction with endomembranes of the Golgi and the endoplasmic reticulum (*20, 27*). *WEE1* did not score as a hit in either of our isogenic synthetic lethal screens (Table S1) or in our analysis of the DepMap data (Fig 1c) indicating that *CCNE1*-amplified cells may have a unique vulnerability to the loss of PKMYT1.

**Fig. 1.**
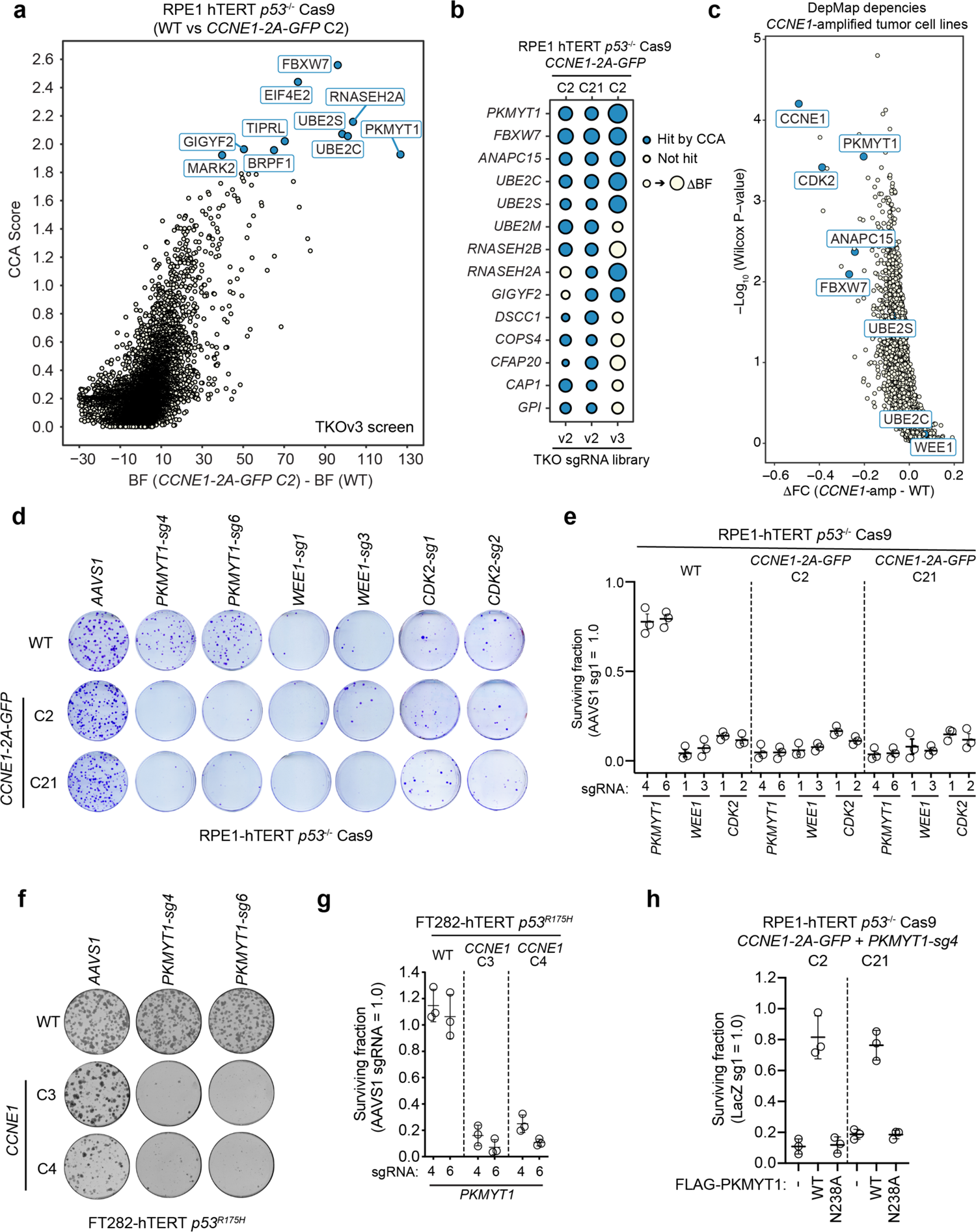
PKMYT1 is synthetic lethal with CCNE1 overexpression. **a**, Synthetic lethal screen performed in RPE1-hTERT *p53^-/-^* Cas9 *CCNE1-2A-GFP* (C2) cells with CCA and ΔBF scores plotted. Gene-level CCA scores and readcounts are available in Table S1 and Datasets S1-S2. **b,** Dot plot of the synthetic lethal hits from three screens performed in RPE1-hTERT *p53^-/-^* Cas9 *CCNE1-2A-GFP* cells. Size of the dots are proportional to ΔBF score and blue colour indicates a hit using CCA (Jenks ranks 4 or 3). **c**, Volcano plot of gene dependencies in cancer cell lines from the DepMap project. Cancer cell lines were grouped according to their CCNE1 amplification status and the difference in median gene depletion was plotted on the x-axis vs the nominal *p*-value of the difference on the y-axis. All values can be found in Table S2. **d,e,** Clonogenic survival assays of the indicated RPE1-hTERT *p53^-/-^ Cas9* cell lines transduced with lentivirus expressing the indicated sgRNAs. Shown in (d) are representative plates with colonies stained with crystal violet. Quantitation is shown in (e). Data are shown as mean ± S.D. (n=3). **f,g,** Clonogenic survival assays of the indicated FT282-hTERT *p53^R175H^* cell lines nucleofected with Cas9 ribonucleoproteins assembled with the indicated sgRNAs. Shown in (f) are representative plates with colonies stained with crystal violet. Quantitation is shown in (g). Data are shown as mean ± S.D. (n=3). **h,** Quantitation of clonogenic survival assays of the indicated RPE1-hTERT *p53^-/-^* Cas9 cell lines transduced with lentivirus expressing *LacZ-sg1* or *PKMYT1-sg4* and doxycycline-inducible sgRNA-resistant *FLAG* alone (-), *FLAG-PKMYT1* or *FLAG*-*PKMYT1-N238A*. Data are normalized to the LacZ-sg1 transduction and shown as mean ± S.D. (n=3).

The identification of a protein kinase, part of a highly druggable enzyme class, prompted us to validate the synthetic lethality between the loss of *PKMYT1* and elevated *CCNE1* levels in the original RPE1 cell line background. We also assessed this genetic interaction in an isogenic set of FT282-based (*28*) *CCNE1*-high hTERT-immortalized fallopian tube cell lines, representing the cell-of-origin of HGSOC (*29*), in which *TP53-R175H* was exogenously introduced. FT282-*CCNE1* cells show accumulation in early S phase, evidence of replication stress, as well as MCM loading defects (Fig S1d-f) (*28*). Two independent sgRNAs targeting *PKMYT1* caused pronounced and selective lethality in the *CCNE1*-high cells in both backgrounds, while sparing their parental counterparts (Fig 1d-g; see Table S3 for TIDE analysis). Reintroduction of an sgRNA-resistant *PKMYT1* transgene fully protected RPE1 *CCNE1*-high cells from depletion of endogenous *PKMYT1*, whereas expression of *PKMYT1-N238A*, which encodes a kinase-dead protein, failed to do so (Fig 1h and Fig. S2a,b). These results indicated that the protein kinase activity of PKMYT1 is essential in cells that are engineered to overexpress cyclin E.

Introduction of sgRNAs targeting *WEE1* and *CDK2* were lethal in RPE1-hTERT cells irrespective of their *CCNE1* status (Fig. 1d-g and Table S3). In contrast, we could generate and stably propagate clonal knockouts of *PKMYT1* in RPE1-hTERT *p53^-/-^* cells that displayed complete loss of CDK1 Thr14 phosphorylation without grossly affecting CDK1 Tyr15 phosphorylation (Fig. S2c,d). Since loss of WEE1 impacts both CDK1 and CDK2 (*26*), and since cyclin E activates CDK2, these observations suggested a simple model where activation of CDK1 was incompatible with viability of cells overexpressing *CCNE1*.

These results prompted us to identify selective inhibitors of PKMYT1 using a structure-guided medicinal chemistry campaign. This work led to the discovery of RP-6306, a highly selective inhibitor of PKMYT1 that has desirable pharmacological properties and is orally bioavailable (Table S4). RP-6306 shows in vitro and cellular selectivity over the related WEE1 kinase (Szychowski et al. in preparation). RP-6306 inhibits PKMYT1 catalytic activity at an IC50 concentration of 3.1 ± 1.2 nM in vitro (Fig. 2a) and has a cellular target engagement EC50 with PKMYT1 of 2.5 ± 0.8 nM using a nanoBRET assay (*30*) whereas its cellular target engagement for WEE1 had an EC50 of 4.8 ± 2.0 μM, or 1920-fold higher (Fig. 2b). RP-6306 treatment led to CDK1, but not CDK2 kinase activation in FT282 *CCNE1* cells, whereas WEE1 inhibition by AZD-1775 led to activation of both kinases, as expected (Fig S3a). In line with PKMYT1 phosphorylating primarily Thr14 on CDK1, in FT282-*CCNE1* cells RP-6306 has an IC50 in reducing CDK1-pT14 of 7.5 ±1.8 nM whereas for CDK1-pY15 the IC50 is over 2 μM (Fig. S3b), a finding which was also confirmed by immunoblotting (Fig. S3c). Both isogenic pairs of *CCNE1*-high cell lines in the RPE1 (Fig 2c and S3d) and FT282 (Fig S3e) backgrounds are sensitive to PKMYT1 inhibition whereas the WEE1 inhibitor (AZD-1775) and two partially selective CDK2 inhibitors (dinaciclib and PF-0687360) did not show consistent *CCNE1* level-dependent sensitivity (Fig 2d and S3d-e). These results are in line with the model that selective CDK1 activation by PKMYT1 inhibition exploits a unique vulnerability of cells with increased *CCNE1* dosage.

**Fig. 2.**
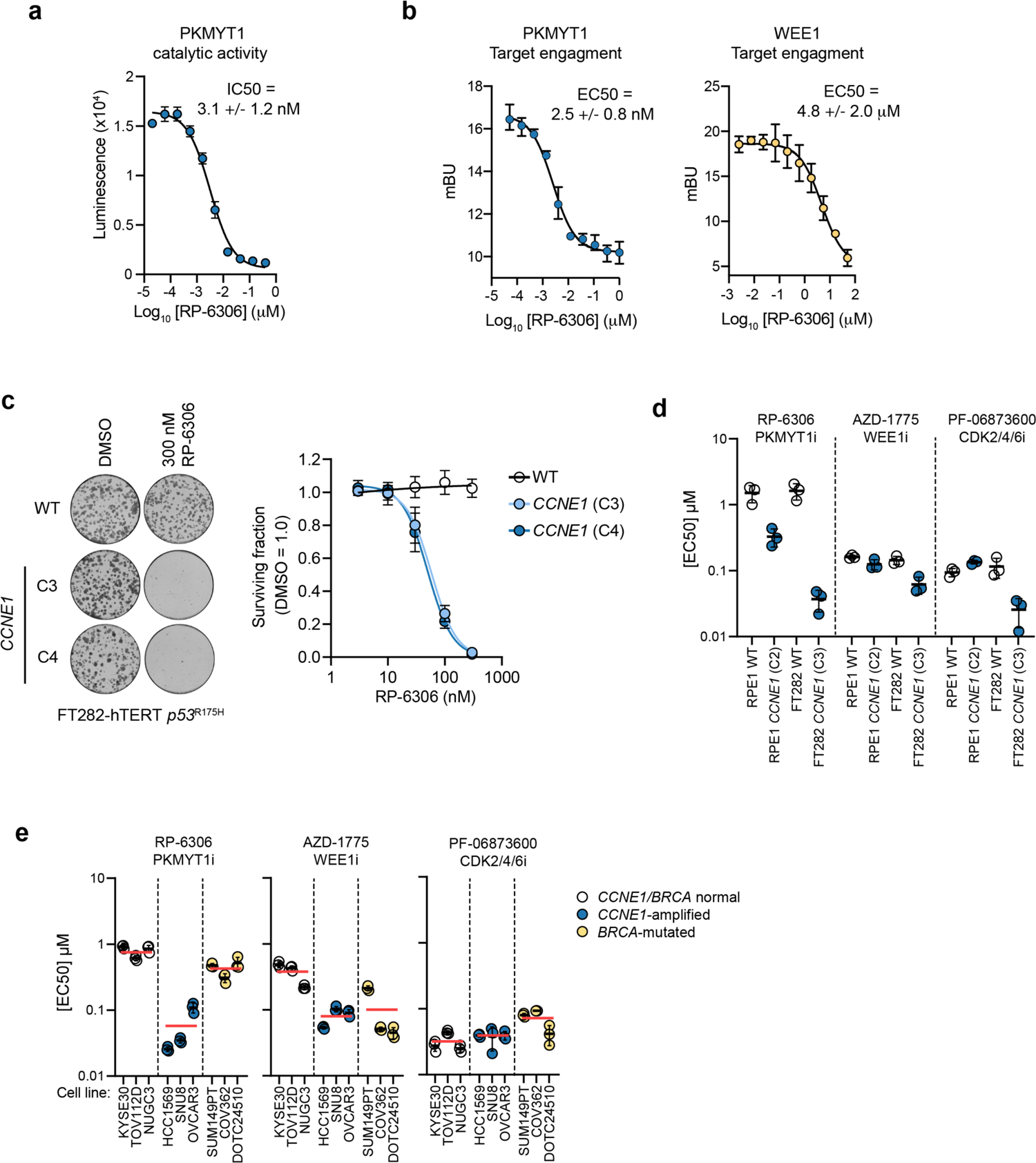
RP-6306 is a selective PKMYT1 inhibitor with activity in CCNE1-amplified cells. **a**, Dose-response of RP-6306 on PKMYT1 catalytic activity as measured with ADP-Glo kinase assay. Data are shown as mean ± S.D. (n=3). **b,** Target engagement of RP-6306 on PKMYT1 (left) and WEE1 (right) using a NanoBRET assay. Data are shown as mean ± S.D. (n=3). **c**, Clonogenic survival of the indicated FT282-hTERT *p53^R175H^* derivatives treated with RP-6306. Shown in left are representative images of plates stained with crystal violet and right is the quantitation. Data are shown as mean ± S.D. (n=3). **d,e**, EC50 determination for growth inhibition for the (d) parental and *CCNE1*-high cells in the RPE1-hTERT *p53 ^-/-^* Cas9 (RPE1), FT282-hTERT *p53^R175H^* (FT282) backgrounds and (e) indicated cancer cell lines treated with the indicated compounds. Growth was monitored with an Incucyte for up to 6 population doublings. Data are shown as mean ± S.D. (n=3). See additional data in Fig. S3d-e. In (e) cell lines are also grouped according to their *CCNE1* or *BRCA* status and the red bar indicates the mean of each grouping.

To test whether PKMYT1 displayed the same selectivity against *CCNE1*-amplification in tumor derived-cell lines, we next assembled a panel of 9 cell lines: 3 with demonstrated amplification or gain of the *CCNE1* locus (HCC1569, SNU8 and OVCAR3), 3 with *BRCA1* or *BRCA2* biallelic mutations that are common in HGSOC (SUM149PT, COV362, DOTC24510), and 3 that are wild type for *CCNE1, BRCA1* and *BRCA2* (KYSE30, TOV112D, NUGC3). For each cell line, we measured the EC50 values for RP-6306, AZD-1775, dinaciclib and PF-0687360 treatment. We found that RP-6306 was, on average, 14.1-fold more cytotoxic to the *CCNE1*-amplified cell lines, with EC50s ranging from 26 to 93 nM, than in cell lines with normal cyclin E levels that yielded EC50s ranging from 616 to 913 nM. In contrast, the WEE1 and CDK2 inhibitors displayed blunted or absent selectivity towards *CCNE1*-amplified cell lines (Fig 2e and Fig S3f). We conclude that PKMYT1 inhibition is selectively cytotoxic to tumor cells displaying *CCNE1* amplification, consistent with the genetic observations made in the isogenic cell lines.

We next explored the basis of the vulnerability of *CCNE1*-high cells to PKMYT1 inhibition using isogenic cell lines. We first assessed whether PKMYT1 inhibition led to DNA damage in *CCNE1*-high cells by monitoring *γ*H2AX levels using quantitative image-based cytometry (QIBC) (*31*). Treatment of these cells with RP-6306 showed that PKMYT1 inhibition led to a dose- and time-dependent accumulation of *γ*H2AX-positive cells solely in the *CCNE1*-high cells in both FT282 and RPE1 backgrounds (Fig 3a and S4a-e). Induction of *γ*H2AX in *CCNE1*-high FT282 cells was recapitulated using sgRNA guides targeting *PKMYT1* (Fig. S4f). Further examination of the *γ*H2AX^+^ population showed that it had DNA content between 1C and 2C but was EdU negative, indicating that cells were not actively replicating DNA (Fig 3a). Imaging of the *γ*H2AX-positive cells by microscopy showed that they had pan-nuclear *γ*H2AX instead of punctate foci and also displayed highly abnormal nuclear morphology, with fragmented or multi-lobed nuclei (Fig 3b). We also observed high levels of micronucleation in FT282 *CCNE1*-high cells treated with RP-6306 (Fig 3c and S4g), consistent with PKMYT1 inhibition causing genome instability in those cells. RP-6306 treatment induced pan-*γ*H2AX in an HCC1569 breast cancer cell line, suggesting that tumor-derived *CCNE1* amplification also renders cells vulnerable to DNA damage induction following PKMYT1 inhibition (Fig 3d and S5a-b).

**Fig. 3.**
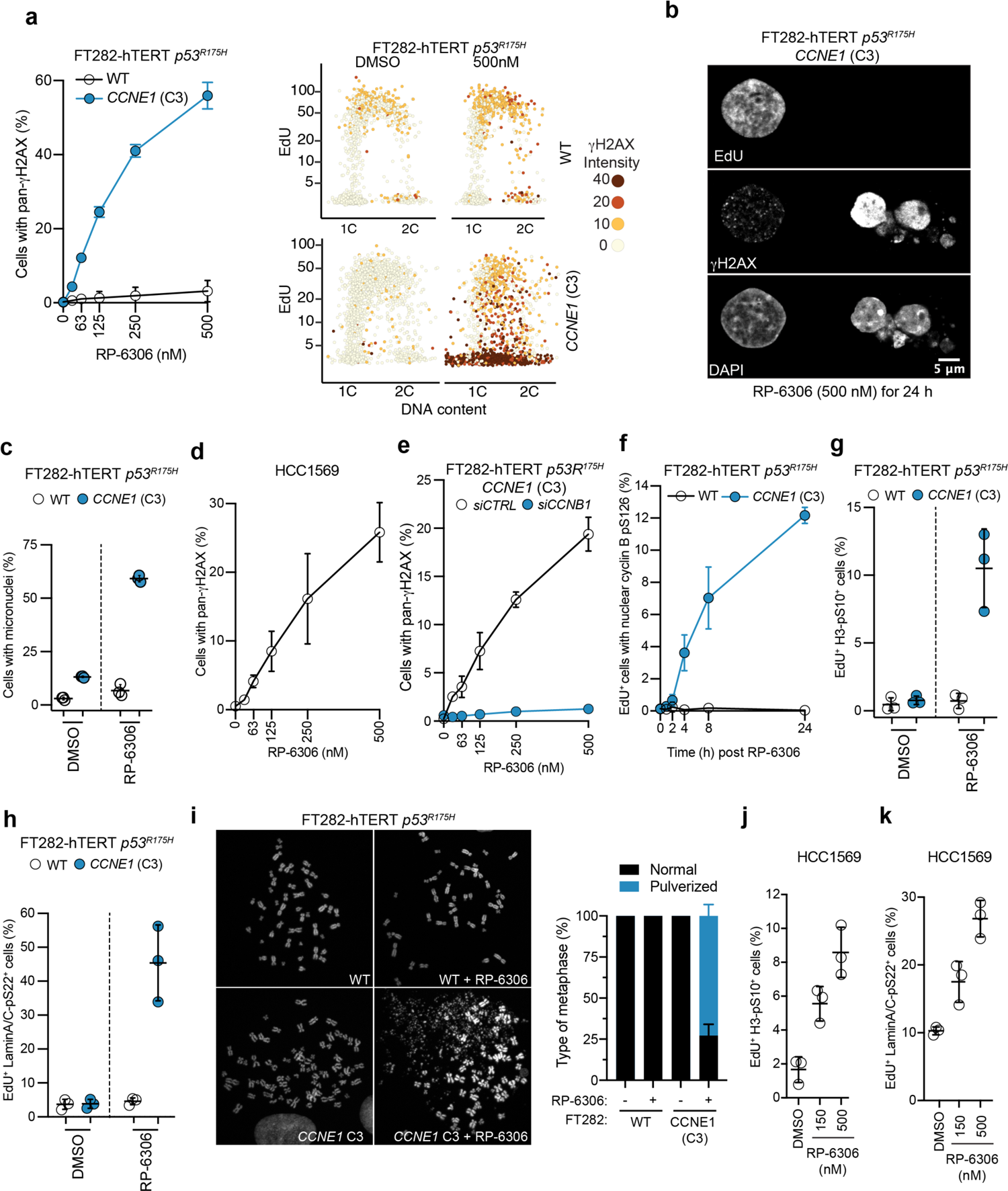
PKMYT1 inhibition causes unscheduled CDK1 activation and mitotic entry in CCNE1-high cells. **a**, QIBC analysis of *γ*H2AX nuclear intensity, EdU incorporation and DNA content (DAPI) in FT282-hTERT *p53^R175H^* cell lines. Representative QIBC plots are shown on the right and on the left is plotted the quantitation of cells with pan-*γ*H2AX as a function of RP-6306 dose. Data are shown as mean ± S.D. (n=3). QIBC validation is shown Fig. S4a. **b**, Representative micrograph showing EdU staining, *γ*H2AX localization and nuclear morphology of FT282 *CCNE1*-high cells. **c**, Quantitation of micronuclei in FT282 parental (WT) and *CCNE1-*high cells following treatment with either RP-6306 (200 nM) or vehicle (DMSO) for 72 h. Data are shown as mean ± S.D. (n=3). Representative micrographs are shown in Fig. S4g. **d**, QIBC quantitation of HCC1569 cells with pan-*γ*H2AX as a function of RP-6306 dose. Data are shown as mean ± S.D. (n=3). QIBC validation is shown Fig. S4g. **e,** Quantitation of FT282-hTERT *p53^R175H^ CCNE1-*high cells transfected with siRNAs targeting cyclin B1 (*siCCNB1)* or non-targeting siRNA (*siCTRL)* with pan-*γ*H2AX as a function of RP-6306 dose. Data are shown as mean ± S.D. (n=3). QIBC validation is shown Fig. S5e. **f**, QIBC quantitation of FT282-hTERT *p53^R175H^* cells of the indicated genotype positive for EdU and cyclin B (CCNB1) pS126 as a function of RP-6306 dose. Data are shown as mean ± S.D. (n=3). QIBC validation is shown Fig. S5h **g**, **h**, Quantitation of double-positive staining for EdU and either histone H3-pS10 (g) or Lamin A/C-pS22 (h) by FACS following vehicle (DMSO) or RP-6306 treatment (500 nM) for 24 h in FT282-hTERT *p53^R175H^* cells of the indicated genotype. Data are shown as mean ± S.D. (n=3). The gating strategy is shown in Fig. S6a. **i**, RP-6306 induces chromosome pulverization. Left, representative micrographs of metaphase spreads of FT282 parental (WT) and *CCNE1-*high cells left untreated or following treatment with either RP-6306 (500 nM) for 24 h. Right, quantitation of cells with pulverized chromosomes. Data are shown as mean ± S.D. (n=3) with at least 60 metaphases counted per replicates. **j**,**k**, Quantitation of double-positive staining for EdU and either histone H3-pS10 (j) or Lamin A/C-pS22 (k) by FACS in HCC1569 cells following vehicle (DMSO) or the indicated dose of RP-6306 for 24 h. Data are shown as mean ± S.D. (n=3). The gating strategy is shown in Fig. S6b.

We next assessed whether the lethality in FT282 *CCNE1*-high cells caused by RP-6306 treatment was due to the activation of CDK1 driven by PKMYT1 inhibition. We introduced CDK1 variants that remove its inhibitory sites on Thr14 or Tyr15 (CDK1-T14A or CDK1-Y15F) or, as a control, maintain a negative charge on these residues (CDK1-T14E/Y15E). We found that introducing CDK1-T14A or CDK1-Y15F profoundly impaired clonogenic potential of *CCNE1*-high cells but not that of their parental counterpart (Fig S5c). Expression of CDK1-TA, which is unable to be phosphorylated by PKMYT1, had the strongest impact on *CCNE1*-high cell viability (Fig. S5c). These data indicate that loss of CDK1 negative regulation by PKMYT1 is toxic in *CCNE1*-high cells.

In parallel experiments, we found that co-treatment of cells with the CDK1 inhibitor RO-3306 abolished RP-6306-dependent pan-*γ*H2AX induction in a dose-dependent manner (Fig S5d). Similarly, depletion of cyclin B1 (*CCNB1*) blocked *γ*H2AX induction, strongly suggesting that the activation of CDK1 by PKMYT1 inhibition is responsible for the DNA damage induction in *CCNE1*-high cells (Fig 3e and S5e-f). *γ*H2AX induction was also dampened by treatment with dinaciclib and PF-06873600, at concentrations that allow for S phase entry, consistent with cyclin E-driven CDK2 activity also being necessary for induction (Fig S5g). However, the lack of selectivity of these inhibitors over other CDKs means that we cannot fully exclude the contribution of other kinases to the phenotype.

Given the role of PKMYT1 in inhibiting cyclin B-CDK1, the mitotic CDK, we posited that the pan-*γ*H2AX terminal phenotype could be secondary to a premature entry into mitosis while cells are still undergoing DNA replication, a phenomenon previously described for WEE1 inhibition (*32, 33*) and in the *Cdk1^AF/AF^* knock-in mouse (*34*). Cyclin B-CDK1 accumulates in the cytoplasm in interphase before its rapid activation that is linked to nuclear translocation and autophosphorylation at the onset of prophase (*35, 36*). We observed that upon RP-6306 treatment, FT282 *CCNE1*-high cells, but not their parental counterpart, accumulate nuclear cyclin B phosphorylated at Ser126 (pS126) in replicating cells marked by EdU incorporation (Fig 3f and S5h-i). Furthermore, following PKMYT1 inhibition, a portion of EdU-positive cells showed evidence of premature entry into mitosis, as measured by histone H3 Ser10 phosphorylation (H3-pS10) and lamin A/C Ser22 phosphorylation (Lamin A/C-pS22; Fig 3g-h, S5j-k and S6a), a marker of nuclear envelope breakdown and lamina disassembly (*37*). RP-6306 treatment of FT282 *CCNE1*-high cells also caused a remarkable increase in chromosome pulverization in M phase-arrested cells, a phenotype that was completely dependent on high *CCNE1* levels (Fig 3i). Chromosome pulverization is a phenotype associated with premature mitotic entry of actively replicating cells (*38*). The lack of premature mitotic entry in the parental FT282 cells following PKMYT1 inhibition is consistent with previous observations that *PKMYT1* depletion is insufficient to trigger premature mitotic entry in HeLa cells (*39, 40*). *CCNE1*-amplified HCC1569 cells also showed dose-dependent increases in H3-pS10 and lamin A/C pS22 signal in replicating cells following RP-6306 treatment, in line with the isogenic cell line data (Fig 3j-k and S6b). Together, these data suggest that the genome integrity in S phase of cells with increased levels of cyclin E is safeguarded by PKMYT1-dependent CDK1 inhibitory phosphorylation.

Cyclin E overexpression causes DNA replication stress that is manifested by premature entry into S phase, unscheduled origin firing, and replication-transcription conflicts (*41*). To test whether replication stress induced by other means also imposes a need for CDK1 inhibitory phosphorylation, we tested whether agents that perturb DNA replication, such as hydroxyurea (HU) or the nucleoside analog gemcitabine, rendered cells sensitive to PKMYT1 inhibition. We observed synergistic cytotoxicity when combining RP-6306 with either HU or gemcitabine in both FT282 parental or FT282 *CCNE1*-high cells, irrespective of cyclin E status (Fig. 4a,b and S7a,b). However, combining gemcitabine (at 0.625 nM) with 62.5 nM of RP-6306 was highly toxic in *CCNE1*-high cells whereas combining the same dose of gemcitabine with higher concentrations of RP-6306 (938 nM) was innocuous in the parental cell line (Fig. 4c). The same trend was observed for combinations of HU and RP-6306 (Fig. S7c) suggesting conservation of a therapeutic index between wild type and *CCNE1*-high cells.

**Fig. 4.**
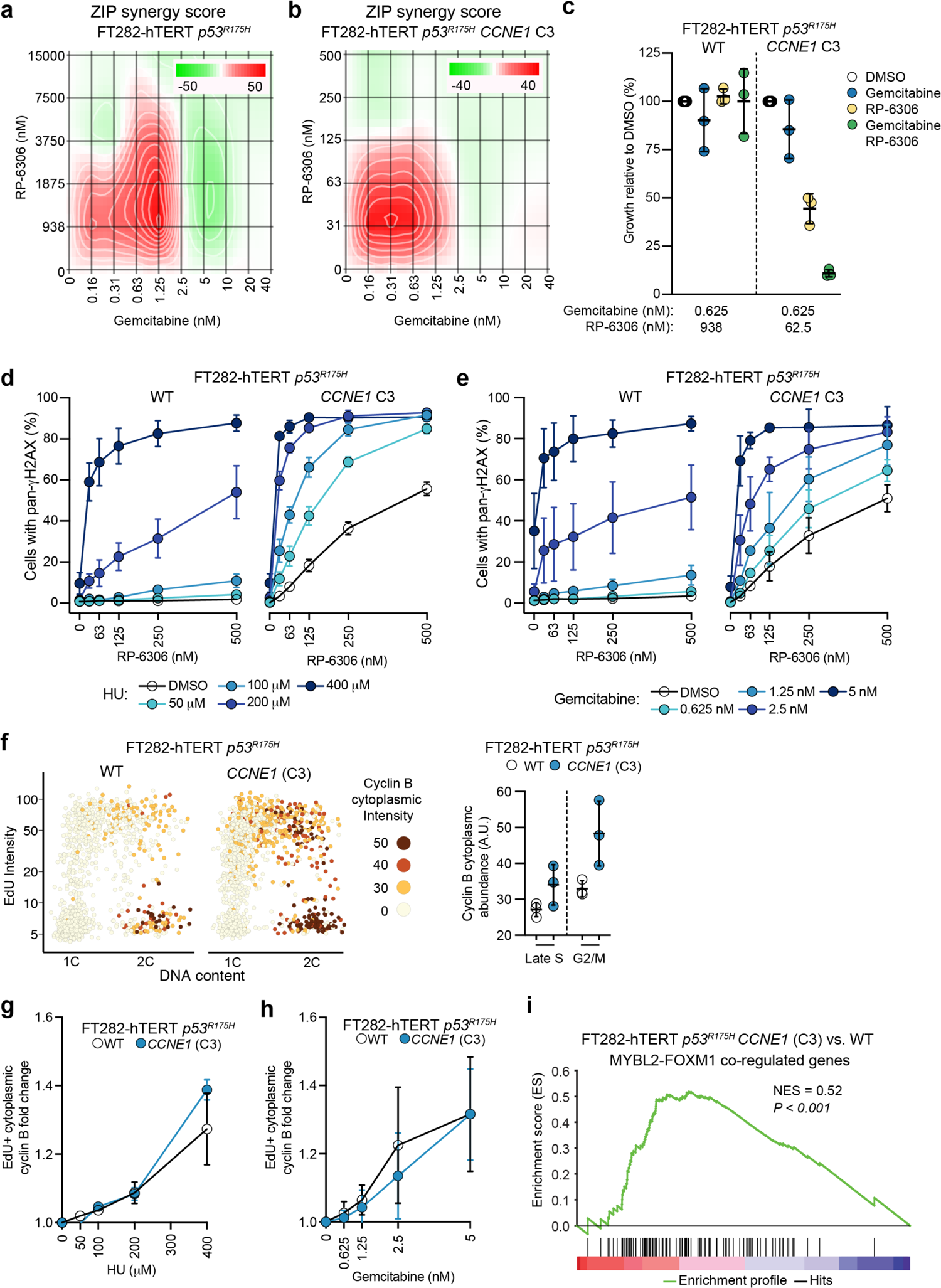
Replication stress and FOXM1-MMB activity underlie vulnerability to PKMYT1 inhibition in CCNE1-high cells. **a,b**, ZIP synergy scores at various dose combinations of RP-6306 and gemcitabine in FT282-hTERT *p53^R175H^* (a) parental and (b) *CCNE1-*high cells. Score ≥10 (red color) represents synergy, ≤-10 (green) represents antagonism. Values were obtained by analyzing mean data from 3 independent biological replicates with SynergyFinder. Growth was monitored with an Incucyte for up to 6 population doublings. **c,** Growth inhibition relative to DMSO control of parental (WT) and *CCNE1-*high cells after treatment with the indicated dose of gemcitabine, RP-6306 or the combination of both. Growth was monitored with an Incucyte for up to 6 population doublings. Data are shown as mean ± S.D. (n=3). **d, e,** QIBC quantitation of cells with pan-*γ*H2AX in response to the indicated RP-6306/HU (d) or RP-6306/gemcitabine (e) combinations. Data are shown as mean ± S.D. (n=3). **f**, QIBC analysis of cyclin B cytoplasmic intensity, EdU incorporation and DNA content (DAPI). Representative QIBC plots are shown on the left and on the right is plotted the quantitation of cells with cytoplasmic cyclin B in late S or G2/M. Data are shown as mean ± S.D. (n=3). QIBC validation is shown Fig. S5i. **g**, **h**, QIBC quantitation of the fold-change of cytoplasmic cyclin B intensity in EdU^+^ cells following treatment with either HU (h) or gemcitabine (i) at the indicated doses for 24 h. Data are shown as mean ± S.D. (n=3). **i**, Gene set enrichment analysis (GSEA) of differential gene expression in FT282 parental (WT) vs *CCNE1*-high (C3) cells for a gene set comprising genes co-regulated by MYBL2 and FOXM1 (Table S5). See additional data in Fig. S8b.

Combining HU or gemcitabine with RP-6306 in FT282 parental cells also caused a synergistic increase in *γ*H2AX signal in replicating (EdU^+^) cells, similar to what was seen in *CCNE1*-high cells (Fig 4d-e). Analogous results were reported with WEE1 inhibition, which also activates CDK1 (*33, 42*). Therefore, these results suggest that the replication stress caused by increased *CCNE1* dosage may be at the root of the vulnerability to unscheduled CDK1 activation.

One interpretation of the above results is that elevated *CCNE1* dosage, or DNA replication stress, renders cells susceptible to unscheduled CDK1 activation and mitotic entry caused by PKMYT1 inhibition. One means by which replication stress could cause this susceptibility is through an increase in CDK1-cyclin B levels or activity. Consistent with this model, we observed that late-S and G2/M cytoplasmic cyclin B levels were elevated in *CCNE1*-high cells, compared to their parental counterparts (Fig 4f and S7d-f). This increase in cyclin B-CDK1 levels was accompanied by higher levels of CDK1 activity in untreated FT282 *CCNE1*-high cells, as measured with immune complex kinase assays (Fig. S7g). The fact that the increase in CDK1 activity was observed only in *CCNE1*-high cells suggests that a consequence of higher cyclin E levels is that CDK1 is primed to become fully activated following PKMYT1 inhibition. Similarly, HU- or gemcitabine-treatment caused an elevation of cyclin B levels in the replicating parental FT282 cells, thereby mimicking the *CCNE1*-high condition (Fig 4g-h). Therefore, the levels and activity of CDK1-cyclin B are elevated in *CCNE1*-high cells, which are likely to make them vulnerable to the loss of CDK1 Thr14 phosphorylation following PKMYT1 inhibition.

The increase in cyclin B levels observed in *CCNE1*-high cells is likely to be at least partly transcriptional, as we and others (*43*) observed that they display an upregulated FOXM1-MYBL2/MuvB (FOXM1-MMB)-dependent transcriptional signature (Fig 4i, S8a,b and Table S5). This correlation between FOXM1 expression and *CCNE1* was also noted in a recent pan-cancer analysis of tumor transcriptomes (*43*). FOXM1 and the MMB complex collaborate to regulate the expression of *CCNB1* and other genes necessary for mitosis (*44*). Therefore, these data suggest that dysregulation of the mitotic transcriptional program by *CCNE1* overexpression could underly the vulnerability of these cells to PKMYT1 inhibition.

Having characterized the impact of PKMYT1 inhibition in vitro, we next assessed whether RP-6306 displayed anti-tumor activity first as a single-agent in tumor xenograft models. This allowed us to explore the pharmacokinetic (PK) and pharmacodynamic (PD) properties of the compound. We implanted *CCNE1*-amplified (HCC1569 and OVCAR3) as well as BRCA1-deficient SUM149PT (Fig S9a) tumor cells as xenografts in mice that were randomised to receive either RP-6306 or vehicle, orally twice daily (BID). We observed dose-dependent tumor growth inhibition (TGI) in both HCC1569 and OVCAR3 cell lines that reached 79 and 84% TGI respectively at 20 mg/kg (Fig 5a,b) whereas RP-6306 had no impact on the growth of SUM149PT-derived tumors at the same dose (Fig S9b), in line with the in vitro data. Mice at each dose level experienced less than 7% body weight loss, suggesting that RP-6306 is well tolerated (Fig S9c,d). We observed a direct dose- and time-dependent relationship between RP-6306 plasma concentration and inhibition of CDK1 Thr14 phosphorylation in tumors (Fig S9e,f), with an EC50 of 2.8 nM (95% CI: 2.0-3.4 nM), indicating very potent on-target activity in vivo. We observed that the levels of cyclin B1 pS126 and histone H3 pS10, as markers of CDK1 activity and M-phase, respectively, are increased 2 h post-administration of RP-6306, suggesting activation of CDK1 and mitotic entry in tumors (Fig S10a,b). We also observed a time-dependent increase of *γ*H2AX following 8 days of treatment with RP-6306 at the 20 mg/kg dose, suggesting that cells with DNA damage accumulate in tumors over time (Fig S10c).

**Fig. 5.**
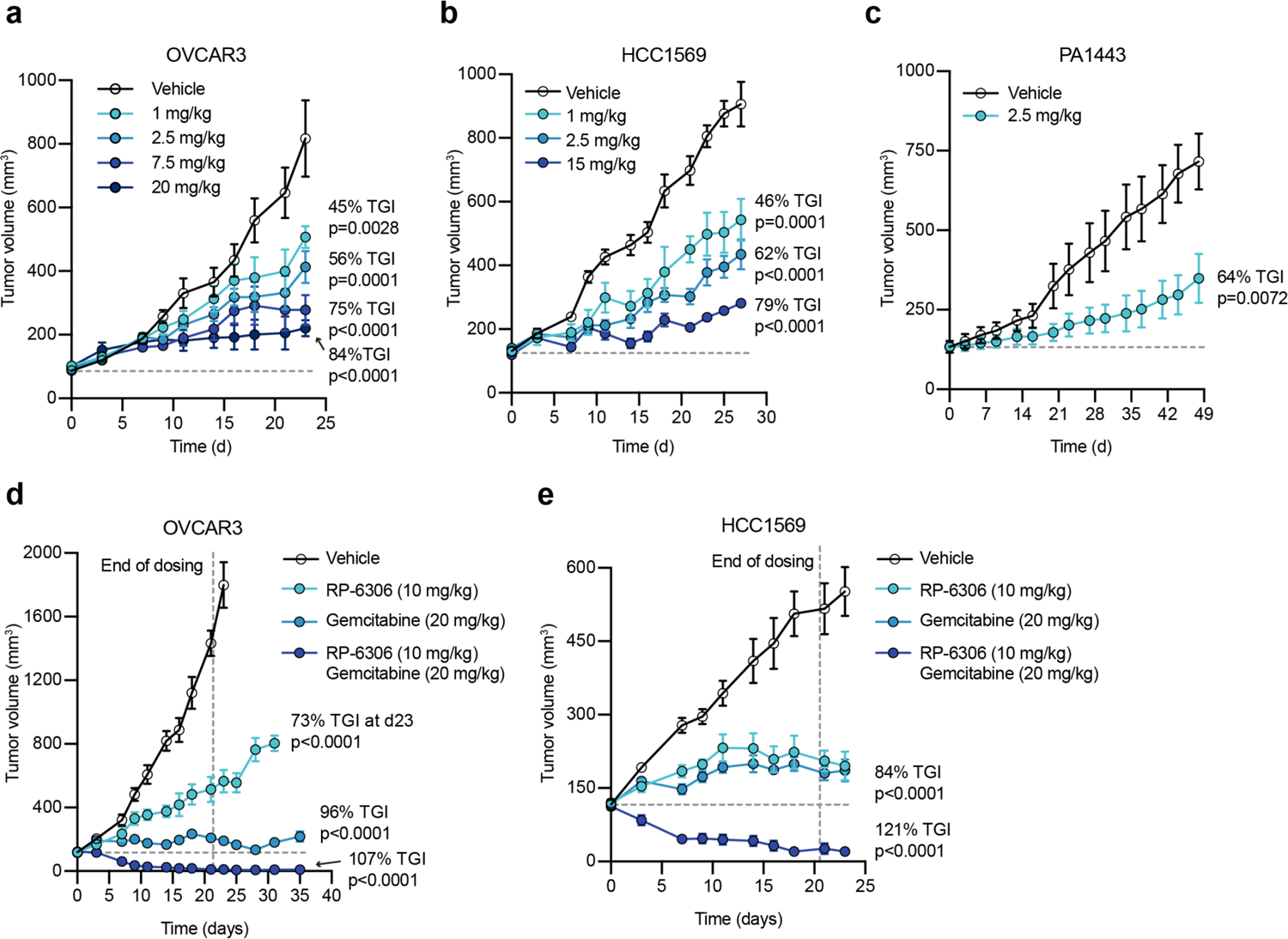
RP-6306 shows single-agent anti-tumor activity and profound tumor regressions in combination with gemcitabine. **a, b**, Tumor growth of OVCAR3 (a) and HCC1569 (b) xenografts in CB-17 SCID and SCID-beige mice treated with either RP-6306 or vehicle. RP-6306 was administered orally BID at the indicated doses for the duration of the experiment. Results are expressed as mean tumor volume ± SEM (n=8). Indicated also is % TGI and *p* values relative to vehicle as determined by a one-way ANOVA. **c**, Tumor growth of a *CCNE1*-amplified pancreatic cancer (PA1443) patient-derived xenograft implanted in BALB/c nude mice treated either with RP-6306 or vehicle. RP-6306 was administered orally BID at 2.5 mg/kg for the duration of the experiment. Results are expressed as mean tumor volume ± SEM (n=8) with % TGI and p-value relative to vehicle as determined by unpaired t-test. **d**, **e**, Tumor growth of OVCAR3 (d) and HCC1569 (e) xenografts in mice treated with either RP-6306, gemcitabine or both. Gemcitabine was administered once weekly intraperitoneally starting at d 0 and RP-6306 was given oral BID for 21 d (e) after which all treatments were stopped, and tumor size monitored for the remainder of the experiment. Results are expressed as tumor volume mean ± SEM (n=7). Indicated also are % TGI and *p-*values relative to vehicle (determined by one-way ANOVA).

The single-agent activity of RP-6306 was also tested in a patient-derived xenograft (PDX) model obtained from a pancreatic adenocarcinoma (model PA1443). This tumor displays moderate *CCNE1* amplification (n=17.5 by whole-exome sequencing; Fig S11a) and increased cyclin E protein levels (Fig S11b-c). PA1443 tumors also harbor *TP53* loss-of-function (G245S) and *KRAS* activating mutations (G12D) typical of this cancer type. BID dosing of RP-6306 at 2.5 mg/kg led to a dose-dependent tumor growth inhibition, reaching 64% over 48 days (Fig 5c) with less than 5% body weight loss (Fig S11d). Together, these data indicate that PKMYT1 inhibition displays single-agent tumor growth inhibition in a variety of *CCNE1*-amplified models.

The observation that replication stress can further sensitize *CCNE1*-high tumor cells to PKMYT1 inhibition prompted us to test whether gemcitabine synergised with RP-6306 in OVCAR3 and HCC1569 tumor xenografts. We selected gemcitabine as it is approved for use in multiple cancer indications, including ovarian cancer, a tumor type where *CCNE1* amplification is prevalent. We tested a dosing regimen where gemcitabine was delivered intraperitoneally once weekly and RP-6306 was given orally BID for 21 days. The gemcitabine/RP-6306 combination led to profound and durable tumor regressions in both OVCAR3 and HCC1569 tumors, with tumors showing no sign of regrowth for up to 15 days following the cessation of the treatment (Fig 5d,e and S12a-d). At day 35, termination of the OVCAR3 model, 3/7 mice were tumor (Fig S12b) and in the HCC1569 tumor group, 2/7 mice remained tumor free at the termination of the experiment on day 30 (Fig S12d). These results indicate a very profound tumor response in both models. A maximum of 10% body weight loss was observed in the combination arm (Fig S12e,f) demonstrating tolerability of the combination. We conclude that enhancing DNA replication stress in *CCNE1*-high tumors may be a particularly effective approach to drive tumor regression in combination with PKMYT1 inhibition.

## Discussion

Oncology drug discovery based on the identification of synthetic lethal interactions holds great promise but very few drug candidates have so far been developed, ab initio, from this approach. In this study, we present how a genome-scale genetic interaction screen in a cellular model of *CCNE1* amplification led to the identification of a vulnerability to PKMYT1 inhibition, and report the discovery of RP-6306, a selective inhibitor of the PKMYT1 kinase that inhibits *CCNE1*-amplified cell and tumor growth. RP-6306 is orally bioavailable and completed studies to support investigational new drug (IND) designation. RP-6306 is expected soon to enter first-in-human clinical studies where its activity against *CCNE1*-amplified tumors as a single agent or in combination will be assessed. This work further demonstrates the applicability of uncovering and prosecuting oncology drug targets via genetic interaction screens.

PKMYT1 is a selective negative regulator of the CDK1 kinase, and we show that the synthetic lethality between PKMYT1 inhibition and the *CCNE1*-high state is caused by unscheduled mitotic entry and lethal genome instability arising from the loss of CDK1 inhibitory phosphorylation. To our surprise, we found that inactivation of the PKMYT1 kinase in cells with normal cyclin E levels is insufficient to induce mitotic entry or wholesale CDK1 activation as determined by cyclin B nuclear translocation or its auto-phosphorylation on Ser126. Similarly, expression of a CDK1-T14A variant was well tolerated in parental FT282 fallopian epithelium cells whereas it was toxic in their *CCNE1*-high counterparts. These observations suggest either that PKMYT1 is minimally active in the S phase of normal cells or that the loss of CDK1 inhibitory phosphorylation on Thr14 in cells without *CCNE1* amplification is buffered by other modulators of CDK1 activity, such as the CDC25 phosphatases, WEE1 activity, CAK kinase regulation or CDK inhibitors (*26*). In contrast, in *CCNE1*-high cells, CDK1 “homeostasis” is impaired causing a vulnerability to the loss of CDK1 inhibitory phosphorylation. This loss of CDK1 homeostasis is likely due to the upregulation of CDK1-cyclin B levels and those of other factors, such as CDC25 phosphatases, through FOXM1-MMB-dependent transcription. The increase in FOXM1-MMB-dependent transcription may be the consequence of replication stress observed in *CCNE1*-high cells, the direct action of cyclin E-CDK2 or both. Together, our observations suggest that the basis for the observed synthetic lethality between PKMYT1 is the result of a one-two punch where the *CCNE1*-driven increase in CDK1-cyclin B in S phase renders cells highly vulnerable to the loss of PKMYT1-driven CDK1 Thr14 inhibitory phosphorylation. The resulting activation of CDK1 causes unscheduled mitotic entry with associated catastrophic genome instability. We note that prior to mitosis, the cyclin B-CDK1 complex accumulates in the cytoplasm and therefore the cytoplasmic PKMYT1 kinase is ideally placed to modulate this latent pool of CDK1-cyclin B.

The one-two punch hypothesis on CDK1 regulation may also explain the noted pan-cellular cytotoxicity of WEE1 loss or inhibition. Unlike PKMYT1, WEE1 also phosphorylates and inhibits CDK2. The resulting activation of CDK2 following WEE1 inhibition causes DNA replication-associated DNA damage and also upregulates CDK1-cyclin B via FOXM1-MMB-dependent transcription. The role of CDK2 in mediating WEE1 cytotoxicity was demonstrated previously in human KMB-7 cells, mouse embryo fibroblasts, and U-2-OS osteosarcoma cells, cell types that do not harbor *CCNE1* amplification (*45, 46*). Therefore, we speculate that the reason why PKMYT1, but not WEE1, shows synthetic lethality with *CCNE1* amplification is due to the selectivity of PKMYT1 for CDK1.

Finally, the elucidation of the mechanism underpinning the synthetic lethality between *CCNE1* overexpression and PKMYT1 suggests avenues for drug combinations that may either drive more durable therapeutic responses or expand patient populations beyond *CCNE1* amplification. Indeed, we show that HU or gemcitabine treatment further enhances cyclin E-driven DNA replication stress leading to profound sensitization of cells and tumors to RP-6306, suggesting that this combination could prove effective in the clinic. These observations also suggest that other agents that perturb DNA replication such as inhibitors of topoisomerase I, PARP or ATR may also display synergy with PKMYT1 inhibition. With respect to additional genetic alterations that could benefit from PKMYT1 inhibition, tumors with mutations in *FBXW7*, which encodes a substrate adaptor for a cullin-RING-ligase that targets cyclin E for ubiquitin-dependent proteolysis (*11, 47–49*), represents an attractive patient population given that cyclin E drives genome instability in these tumors (*50*). We finally note that alterations in FOXM1-MMB-driven transcription are seen in multiple cancers (*51–53*), including HPV-positive head and neck squamous cell carcinoma, where it causes sensitivity to WEE1 inhibition (*54*). Therefore, testing whether FOXM1-MMB-driven transcription can act as a more general vulnerability to the loss of CDK1 inhibitory phosphorylation may further expand the range of tumors that could benefit from PKMYT1 inhibitors.

## Methods

### Cell lines and cell culture

All cell lines were grown at 37°C and 5% CO_2._ RPE1-hTERT *p53^-/-^* Cas9 and RPE1-hTERT *p53^-/-^* Cas9 *PKMYT1^-/-^* cells were grown in DMEM (Life technologies # 11965-092) with 10% FBS (Wisent #080150) and 1% Pen/Strep (Wisent #450-201-EL). RPE1-hTERT *p53^-/-^* Cas9 CCNE1 overexpressing cell lines were constructed by piggyBac transposition of *CCNE1-2A-GFP* into the parental cell line and selection of clones with mid-range GFP expression. FT282-hTERT *p53R^175H^* WT (empty vector) and CCNE1 overexpressing cell lines were obtained from Ronny Drapkin (*28*) and cultured in DMEM: F-12(1:1) (Life technologies # 11330-032) with 5%FBS, 1% UltroserG (Pall Life Sciences #15950-017) and 1% Pen/Strep. 293T cells were cultured in DMEM with 10% FBS and 1% Pen/Strep. HEK293T cells were cultured in DMEM with 10% FBS and 1% Pen/Strep.. HCC1569 cells were cultured in RPMI 1640 (Life technologies # 118575-093) with 10% FBS and 1% Pen/Strep. SNU8 cells were cultured in RPMI 1640 with 10% FBS, 1% Pen/Strep, 25mM HEPES. OVCAR3 cells were cultured in RPMI 1640 with 20% FBS, 1% Pen/Strep and 0.01mg/mL insulin. SUM149PT cells were cultured in Ham’s F12 (Life technologies # 11765-054) with 5% FBS, 10mM HEPES, 1% Pen/Strep, 1ug/mL hydrocortisone and 5ug/mL inulin. KYSE30 cells were cultured in 45% RPMI 1640 with 45% Ham’s F12, 10% heat inactivated. FBS and 1% Pen/Strep. TOV112D cells were cultured in 42.5% MCDB 105, 42.5% Medium 199 (Life technologies #11150-059), 15% FBS and 1% Pen/Strep. NUGC3 cells were cultured in RPMI 1640 with 10% FBS and 1% Pen/Strep. COV362 cells were cultured in DMEM with 10% FBS and 1% Pen/Strep. DOTC23510 cells were cultured in DMEM with 10% FBS and 1% Pen/Strep. The OVCAR3 and HCC1569 cells were shown to have amplified *CCNE1* in (*55, 56*) whereas SNU8 was shown to have *CCNE1* copy number gain in the CCLE database (https://portals.broadinstitute.org/ccle).

## Plasmids

For CRISPR-Cas9 genome editing, sgRNAs were cloned either in lentiCRISPRv2 or in lentiguide-NLS-GFP as in (*57*). For PKMYT1 overexpression in cells, an N-terminally 3xFLAG-tagged PKMYT1 open reading frame (CCDS10486.1) was cloned into the pDONR221 Gateway entry vector (Thermo Fisher Scientific, 12536017). Mutagenesis was performed by PCR to generate a PKMYT1 sgRNA-resistant version carrying silent mutations between nucleotides 966 and 981 (tgagttcactgccggt to cgaatttaccgctggc) and the kinase-dead mutant N238A. PKMYT1 coding sequences were transferred by Gateway technology to the destination vector pCW57.1 (Addgene: 41393) used for transduction in cells. For CDK1 mutant expression in cells the coding sequence for CDK1-TY14,15AF-GFP was gene synthesized and cloned into the pHIV-NAT-hCD52 vector (a kind gift from Ralph Scully) using EcoRI and BamHI restriction enzymes. Mutagenesis was then performed to revert each phosphosite back to the wild type amino acids to create CDK1-T14A-GFP and CDK1-Y15F-GFP. Phospho-mimetic mutants were generated by mutating CDK1-TY14,15AF-GFP to CDK1-TY14,15EE. The sgRNA sequences used in this study are included in Table S6.

### Lentiviral transduction

Lentiviral particles were produced in 293T cells in 10-cm plates by co-transfection of 10 μg of targeting vector with 3 μg VSV-G, 5 μg pMDLg/RRE and 2.5 μg pRSV-REV (Addgene #14888, #12251, #12253) using calcium phosphate. Medium was refreshed 12–16 h later. Virus-containing supernatant was collected ∼36-40 h post transfection and cleared through a 0.2-μm filter. Viral transductions were performed in the presence of polybrene (Sigma-Aldrich, 4 μg/mL RPE1-hTERT *p53^-/-^* Cas9 and 16 μg/mL FT282-hTERT *p53-R175H*) at a multiplicity of infection (MOI) <1.

### Antibodies

Primary antibodies used in this study include: histone H2A.X (phospho-S139, Cell Signalling Technologies #2577, 1:500 IF), histone H2A.X (phospho-S139, Millipore Sigma #05-636, 1:500 IF), CDK1 (Thermo Fisher Scientific #33-1800, 1:1000 IB and ELISA), CDK1-phosphoT14 (Abcam #ab58509, 1:1000 IB and ELISA), CDK1-phoshoY15 (Cell Signaling Technology #9111, 1:1000 IB), PKMYT1 (Bethyl A302-424A, IB), Histone H3-phosphoS10 (Cell Signaling Technology #9706, 1:500 FC), lamin A/C (Cell Signaling Technology 4C11 #4777, 1:500 IF), lamin A/C-phosphoS22 (Cell Signaling Technology D2B2E #13448, 1:500 FC and IF), Cyclin B1 (Cell Signalling Technologies #2577, 1:500 IF, 1:1000 IB), *α*-Tubulin (Millipore DM1A CP06, 1:4000 IB), CDK2 (Upstate 05-596, 1:1000 IB), Cyclin B1-phosphoS126 (Abcam ab55184, 1:500 IF), MCM2 (BD Biosciences 610700, 1:250 IF), MCM4 (Novus Bioogicals H0004137-B01P, 1:500 IF), CHK1-phosphoS345 (Bethyl 2348, IB), Cyclin E1 (Abcam ab3927 IB or Cell Marque #AC0120RUO IHC) Alpha Actinin (Millipore Sigma 05-384, IB), Vinculin (Cell Signalling 13901S, 1:1000 IB). The following agarose-coupled antibodies were used for immunoprecipitation (IP) in kinase assays: CDK1 (Santa Cruz sc-54 AC) and CDK2 (Santa Cruz sc-6248 AC). The following secondary antibodies were used for immunoblotting (IB): anti-mouse Irdye 800CW, anti-rabbit IRdye 680RD (926-32210 and 926-68071; LiCOR, 1:5000), anti-Mouse IgG HRP (Cedarlane # NA931-1ML, 1:4000), anti-Rabbit IgG HRP (Cedarlane # 111-035-144, 1:4000), anti-Rabbit IgG HRP (abcam 97051, 1:10,000). The secondary antibody used for ELISA was anti-Rabbit IgG HRP (Jackson Immunoresearch #111-035-144). The following secondary antibodies were used for immunofluorescence (IF) and flow cytometry (FC): AlexaFluor488 donkey anti-rat IgG (Thermo Fisher Scientific A21208, 1:1000), AlexaFluor647 donkey anti-mouse IgG (Thermo Fisher Scientific A31571, 1:1000), AlexaFluor488 goat anti-mouse IgG (Thermo Fisher Scientific A11029, 1:1000), AlexaFluor647 goat anti-rabbit IgG (Thermo Fisher Scientific A21244, 1:1000). Finally, the following secondary antibodies were used for AlphaLISA assays: AlphaLISA anti-rabbit IgG Acceptor beads (Perkin Elmer #AL104C) and AlphaLISA anti-mouse IgG Donor beads (Perkin Elmer #AS104D)

### Short interfering RNAs

siRNA oligonucleotides (siCTRL ON-TARGET Plus D-001210-03-50 and siCCNB1 ON-TARGET Plus L-003206-00-0005; Dharmacon) were transfected in Opti-MEM reduced-serum medium using Lipofectamine RNAiMAX agent (Thermo Fisher Scientific #13778-075) following manufacturer recommended protocol. Fresh media was added to cells 16 h after transfection. Cells were used for high content imaging and immunoblotting 48 h after transfection.

### Fine chemicals

The following drugs were used in the course of the study: RP-6306 (this study), AZD1775 (Selleckchem, S1525), dinaciclib (MedChemExpress, HY-10492), PF-06873600 (MedChemExpress, HY-114177), RO-3306 (Selleckchem, S7747), gemcitabine (Cayman Chemicals, 9003096) and hydroxyurea (Sigma-Aldrich #H8627). Concentration and duration of treatment is indicated in the legends of the corresponding figures.

### CRISPR screens

CCNE1-overexpression synthetic lethality screens were conducted as three parallel screens with a parental cell line and two isogenic clones overexpressing *CCNE1* (C2 and C21). For the screens, RPE1-hTERT Cas9 *p53*^-/-^ parental and RPE1-hTERT Cas9 *p53^-/-^ CCNE1*-overexpressing clones were transduced with the lentiviral TKOv2 sgRNA library at a low MOI (∼0.3) and media containing 20 μg/mL puromycin (Life Technologies) was added the next day to select for transductants. The following day, cells were trypsinized and replated in the same plates while maintaining puromycin selection. 3 d after infection, which was considered the initial time point (t0), cells were pooled together and divided into 2 sets of technical replicates. Cells were grown for a period of 18 d and cell pellets were collected every 3d. Each screen was performed as a technical duplicate with a theoretical library coverage of ≥ 400 cells per sgRNA maintained at every step. Genomic DNA was isolated using the QIAamp Blood Maxi Kit (Qiagen) and genome-integrated sgRNA sequences were amplified by PCR using NEBNext Ultra II Q5 Master Mix (New England Biolabs). i5 and i7 multiplexing barcodes were added in a second round of PCR and final gel-purified products were sequenced on an Illumina NextSeq500 system at the LTRI NBCC facility (https://nbcc.lunenfeld.ca/) to determine sgRNA representation in each sample. Later, another screen was conducting using the next-generation TKOv3 library in RPE1-hTERT Cas9 *p53*^-/-^ parental and -*CCNE1* (C2) cells using the same procedure outlined above.

### DepMap data mining

CRISPR dependency data (CERES scores) (*18, 58*) and gene-level copy number data (*59*) were downloaded from the 2021 Q1 DepMap release using the Broad Institute’s DepMap portal. Cell lines were characterized as being “*CCNE1*-amplified” if they had a copy number value that was greater than 1.58 (approximately equal to 2x total copy number relative to ploidy), or “WT” if they had a copy number value that was less than or equal to 1.58; cell lines with no copy number data for *CCNE1* were removed from the analysis. From a total of 808 cell lines in the dependency data set, 6 were removed, 20 were classified as “*CCNE1*-amplified,” and 782 were classified as “WT.” The Wilcoxon Rank Sum Test was used to compare dependency scores for each gene between the 2 groups. Nominal *p*-values are provided. Results of the analysis can be found in a tabular format in Supplementary Table 2.

### Clonogenic survival assays

Cells were seeded in 6-well plates, 300 cells per well for RPE1 and 400 for FT282. Single cells were grown out until distinct colonies formed with greater than 50 cells per colony. Colonies were rinsed with PBS and stained with 0.4% (w/v) crystal violet in 20% (v/v) methanol for 30 min. The stain was aspirated, and plates were rinsed twice in double-distilled H_2_O and air-dried. Colonies were counted using a GelCount instrument (Oxford Optronix, GelCount).

### Cell proliferation assays

RPE1-hTERT Cas9 *p53^-/-^*, FT282-hTERT *p53^R175H^* and their respective *CCNE1*-overexpressing isogenic pairs were seeded in 96-well plates (Corning Costar #5595) at a density of 150 cells per well for RPE1-hTERT Cas9 *p53*^-/-^ *CCNE1* (C2) or 100 cells per well for all others. After 24 h, cells were treated using an automated D300e digital dispenser (Tecan) at drug concentrations ranging from 0.15 nM to 3 µM. Medium and drugs were refreshed every 3-4 days and cellular confluency was monitored up to 6 population doublings using an IncuCyte S3 Live-Cell Imager (Sartorius). Percent confluence relative to a non-treated control was used to evaluate growth inhibition induced by test compounds. Synergy between RP-6306 and hydroxyurea (HU) or gemcitabine was analyzed using the online SynergyFinder tool (*60*) using the ZIP model (*61*) (https://synergyfinder.fimm.fi).

### Immunofluorescence

Cell were seeded onto glass coverslips and treated as indicated in the figure legends. Prior to harvesting, cells were pulsed with 20 μM EdU (5-ethynyl-2-deoxyuridine, Life Technologies #A10044) for 30 min and then washed with PBS and fixed with 4% paraformaldehyde (PFA) in PBS for 15 min at room temperature (RT). Cells were then rinsed with PBS and permeabilized using 0.3% Triton X-100/ PBS for 30 min. For chromatin-bound MCM measurements, cells were pre-extracted for 15 min on ice with CSK buffer (300 mM sucrose, 100 mM NaCl, 3 mM MgCl_2_, 10 mM PIPES pH 7.0, 0.5% v/v Triton-X 100) before PFA fixation. Cells were washed with PBS and incubated in blocking buffer (10% goat serum (Sigma #G6767), 0.5% NP-40 (Sigma-Aldrich, #I3021), 5% w/v Saponin (Sigma-Aldrich, #84510), diluted in PBS) for 30 min. Fresh blocking buffer containing primary antibodies was added for 2 h. Cells were rinsed three times with PBS and then blocking buffer, with secondary antibodies and 0.4 μg/mL DAPI (4,6-diamidino-2-phenylindole, Sigma-Aldrich, #D9542) was added for 1 h. After rinsing with PBS, immunocomplexes were fixed again using 4% PFA/PBS for 5 min. Cells were rinsed with PBS and incubated with EdU staining buffer (150 mM Tris-Cl pH 8.8, 1mM CuSO_4_, 100 mM ascorbic acid and 10 μM AlexaFluor 555 azide (Life Technologies, #A20012) for 30 min. After rinsing with PBS coverslips were mounted onto glass slides with ProLong Gold mounting reagent (Invitrogen, #P36930) Images were acquired using a Zeiss LSM780 laser-scanning microscope (Oberkochen, Germany).

### High content imaging and quantitative image-based cytometry (QIBC)

For high-throughput analysis of nuclear γ-H2AX, cells were seeded in 96-well plates (3000 cells/well for FT282-hTERT *p53^R175H^*) and cultured for up to 72 h depending on the experiment. Cells were fixed, permeabilized and stained in the same manner as immunofluorescence described above. Wells were filled with 200 μl PBS and images were acquired at the Network Biology Collaborative Centre (LTRI) on an InCell Analyzer 6000 automated microscope (GE Life Sciences) with a 20X objective. Image analysis was performed using Cellprofiler (Fig S13) (*62*).

### Immunoblotting

Cell pellets were extracted by incubation in NP-40 lysis buffer (50 mM Tris-Cl pH 7.4, 250 mM NaCl, 5 mM EDTA, 1% NP-40, 0.02%, NaN_3,_ 1x protease inhibitor cocktail (Roche, #11836170001)),for 30 min on ice. Extracts were cleared by centrifugation at 13,000x *g* for 10 min at 4°C. Cleared extracts were diluted in 2x sample buffer (20% glycerol, 2% SDS, 0.01% bromophenol blue, 167 mM Tris-Cl pH 6.8, 20 mM DTT) and boiled prior to separation by SDS-PAGE on Novex Tris-glycine gradient gels (Invitrogen, #XV0412PK20). Alternatively, cell pellets were boiled directly in 2x sample buffer before separation by SDS-PAGE. Proteins were transferred to nitrocellulose membranes (VWR, #CA10061-152), then blocked with in 5% milk TBST and probed overnight with primary antibodies. Membranes were washed three times for five minutes with TBST, then probed with appropriate secondary antibodies for one hour, and washed again with TBST, three times for five minutes. Secondary antibody detection was achieved using an Odyssey Scanner (LiCOR) or enhanced chemiluminescence (ECL SuperSignal West Pico, Thermo Fisher Scientific #34579).

### Flow cytometry

Cells were pulsed with 20 μM EdU (5-ethynyl-2-deoxyuridine, Life Technologies #A10044) for 30 min, collected by trypsinization, resuspended as single cells, washed 1 x in PBS and pelleted at 600x *g* for 3 min at 4°C. All subsequent centrifugations were performed in this manner. Cells were fixed in 4% PFA/PBS for 15 min at RT, excess ice cold PBSB (1% BSA in PBS, 0.2 μM filtered) was added before pelleting. Cells were resuspended in permeabilization buffer (PBSB, 0.5% Triton-X 100) and incubated at RT for 15 min. Excess blocking buffer (PBSB, 0.1% NP-40) was added, cells were pelleted, resuspended in blocking buffer containing primary antibodies and incubated at RT for 1 h. Excess blocking buffer containing secondary antibodies was added, cells were pelleted, resuspended in blocking buffer and incubated at RT for 30 min. Excess blocking buffer was added, cells were pelleted and washed one additional time in PBSB. Cells were resuspended in EdU staining buffer (150 mM Tris-Cl pH 8.8, 1mM CuSO_4_, 100 mM ascorbic acid and 10 μM AlexaFluor 555 azide (Life Technologies, #A20012)) and incubated at RT for 30 min. Excess PBSB was added, cells were pelleted and washed one additional time in PBSB. Cell were resuspended in analysis buffer (PBSB, 0.5 µg/ml DAPI, 250 µg/µl RNase A (Sigma-Aldrich, #R4875)) and incubated at 37°C for 30 min or left at 4°C overnight. Cells were analyzed on a Fortessa X-20 (Becton Dickinson) with at least 10,000 events collected and analyzed using FlowJo v10.

### Immune complex histone H1 kinase assays

Cell pellets were resuspended in 250 μL EBN buffer (150 mM NaCl, 0.5% NP-40, 80 mM *β*-glycerol phosphate (Sigma-Aldrich, #50020), 15 mM MgCl_2,_ 20 mM EGTA, 1 mg/mL ovalbumin (Sigma-Aldrich, #5503), 1x protease inhibitor cocktail (Roche, #11836170001), pH 7.3) and incubated on ice for 5 min. Cell lysis was induced by two freeze-thaw cycles of incubation in liquid nitrogen and a 37°C water bath, and lysates were cleared by centrifugation at 13000x *g* at 4°C for 10 min. Protein concentration was determined by Bradford assay (Thermo Fisher Scientific #1856209). For immunoprecipitation of kinases, 200 μg of extract was diluted in 750 μL EBN buffer and 10 μg of CDK1 or CDK2 primary antibody agarose bead conjugates were added to the extract and rotated at 4°C overnight. Immunoprecipitates were pelleted by centrifugation at 2500x *g* at 4°C for 5 min and washed 2x in 750 μL EBN followed by 1 mL EB (80 mM *β*-glycerol phosphate, 15 mM MgCl_2,_ 20 mM EGTA, 1 mg/mL ovalbumin). After the final wash, the immunoprecipitates were resuspended in 500 μL EB and split into 2x samples. One sample was used for immunoblot analysis and the other was carried forward for kinase assays. Following removal of the final wash, immunoprecipitates were resuspended in 11 μl histone H1 kinase assay buffer (80 mM *β*-glycerol phosphate, 15 mM MgCl_2,_ 20 mM EGTA, 1 mg/mL ovalbumin, 10 mM DTT, 0.15 μg/μL Histone H1 (Sigma-Aldrich, #H1917), 22 μM ATP, 0.05 μCi/μL *γ*^32^P-ATP (Perkin Elmer NEG502A250UC), pH 7.3) and incubated at room temperature for 30 min. Reactions were quenched by addition of 5 μL 6x sample buffer (60% glycerol, 6% SDS, 0.03% bromophenol blue, 1500 mM Tris-Cl pH 6.8, 60 mM DTT) and resolved by SDS-PAGE. Gels were exposed to a phosphor imaging screen for 1-2 d and imaged using a Typhoon FLA 9500 (GE Healthcare Life Sciences).

### Cytogenetic analyses

1.5 x 10^6^ FT282-hTERT *p53^R175H^* cells were seeded in 10-cm dishes. 24 h later RP-6306 was added at a final concentration of 500 nM for 24 h. 100 ng/mL KaryoMAX colcemid (Thermo Fisher Scientific #15212-012) was added to the media in the last 2 h of treatment and cells were harvested as follows: Growth medium was stored in a conical tube. Cells were treated twice for 5 min with 1 mL of trypsin. The growth medium and the 2 mL of trypsinization incubations were centrifuged (250x *g* 5 min, 4°C). Cells were then washed with PBS and resuspended in 75 mM KCl for 15 min at 37°C. Cells were centrifuged again, the supernatant was removed, and cells were fixed by drop-wise addition of 1 mL fixative (ice-cold methanol: acetic acid, 3:1) while gently vortexing. An additional 9 mL of fixative was then added, and cells were fixed at 4°C for at least 16 h. Once fixed, metaphases were dropped on glass slides and air-dried overnight. To visualize mitotic cells, slides were mounted in DAPI-containing ProLong Gold mounting medium (Invitrogen, #P36930). Images were captured on a Zeiss LSM780 laser-scanning confocal microscope

### RNA-seq sample preparation, sequencing and analysis

Cells were seeded in 10-cm dishes (2.5 x 10^6^ FT282-hTERT *p53^R175H^* WT or 2 x 10^6^ *CCNE1* (C3 and C4)). The next day, cells were collected by trypsinization, washed once in PBS, and then pelleted. Pellets were snap-frozen in liquid nitrogen. RNA extraction and sequencing of the full transcriptome was performed using NovaSeq at BGI Hong Kong. Raw FASTQ files from a paired-end library were assessed using the FastQC v0.11.9 software (http://www.bioinformatics.babraham.ac.uk/projects/fastqc/) to determine the quality of the reads; read length was 150bp. The FASTQ files were then aligned to the GENCODE GRCh38 v36 primary assembly of the human genome and quantified using Salmon v1.4.0 (*63*) with the command line flags “--validateMappings” and “--gcBias” to obtain read counts. Raw counts were processed using the bioconductor package edgeR v3.30.3 in R (*64*). Genes expressed with counts per million (CPM) > 0.1 in at least two samples were considered and normalized using trimmed mean of M-values (TMM) to remove the library-specific artefacts. For subsequent analyses, voomY transformation was applied to RNA-seq count data to obtain normalized expression values on the log2 scale. Heatmaps were generated using the package heatmap3 v1.1.9 in R. Unsupervised hierarchical clustering was performed by calculating distances using the Pearson correlation metric and clustering using the complete method. Gene expression values were averaged and scaled across the row to indicate the number of standard deviations above (red) or below (blue) the mean, denoted as row z-score. Gene set enrichment analysis (GSEA) (*65*) was performed to identify the enrichment of genes co-regulated by MYBL2-FOXM1 (*66*) in the FT282-hTERT *p53^R175H^ CCNE1* C3 and C4 clones compared to parental WT cells. Analysis was performed using 1000 permutations of the gene set, and normalized enrichment scores (NES) were obtained to reflect the degree to which the gene set is overrepresented in the FT282-hTERT *p53^R175H^ CCNE1* C3 and C4 clones.

### ADP-GLO assay

For the ADP-Glo assay (Promega, Cat #V9103) human recombinant PKMYT1 (full-length human GST-PKMYT1 recombinant protein; Thermo Fisher Scientific A33387, Lot 1938686), was diluted in enzyme assay buffer (70 mM HEPES, 3 mM MgCl_2_, 3 mM MnCl_2_, 50 mg/mL PEG20000, 3 mM Sodium-orthovanadate, 1.2 mM DTT) in a 5 mL volume and plated in 384-well plates (to a final concentration of 18.5nM) followed by addition of 5 μL enzyme assay buffer. The enzyme/compound mix was incubated at room temperature for 15 min before addition of 5 μL 30 mM ATP (diluted in enzyme assay buffer) so that the final ATP concentration was 10 mM. After incubation at 30°C for 1 h, 15 μL of ADP-Glo reagent was added and incubated at room temperature for 40 min. Finally, 30 μL of the kinase detection reagent was added, the plate was incubated at RT for 30 min and luminescence was measured using an EnVision plate reader (Perkin-Elmer). Luminescence is measured for 0.25 seconds/well and rate per second was obtained by multiplying the luminescence value by 4.

### NanoBRET assay

To determine the affinity of RP-6306 in the PKMYT1 or WEE1 NanoBRET target engagement assay (Promega Corp.), HEK293T cells were transfected with a NanoLuc fusion vector for PKMYT1 (Promega NV1871) or WEE1 (Promega NV2231) with transfection carrier DNA (Promega E4881) using Fugene HD Transfection reagent (Promega E2311) in Opti-MEM without phenol red (Thermo Fisher Scientific, 11058021) and incubated overnight. Cells were trypsinized, counted and 17000 cells/well were plated in 96-well plates with K-5 cell-permeable kinase NanoBRET TE tracer (Promega N2482) and RP-6306 and incubated for 2 h at 37°C. Intracellular TE Nano-Glo Substrate/Inhibitor (Promega N2160) was added, and the intensity of the acceptor emission (610 nm) and the donor emission (450 nm) were measured using an EnVison plate reader (Perkin-Elmer)

### AlphaLISA assay

HCC1569 cells were plated into a 96-well TC-treated culture plate (30000 cells/well) and grown overnight. The next day, RP-6306 was dispensed using a Tecan D300e digital dispenser with 3-fold dilutions. After compound addition, cell plates were centrifuged at 300x *g* for 10 seconds, and then placed in the incubator for 2 hours. Cells were lysed in AlphaLISA lysis buffer supplemented with 1X protease (Roche, #11836170001), and phosphatase inhibitors (Sigma-Aldrich #4906837001) and 1 mM PMSF. Plates were rotated at 500x *g* for 20 min to facilitate lysis. Plates were then sealed with aluminum foil and frozen at −80°C for at least 1 hour. Lysates were thawed at 37°C for 10 min and 10 μl of each lysate was transferred in duplicate to 384 well assay plates. Antibodies were added at a final concentration of 5 nM or 1 nM for CDK1-pT14/CDK1-total or CDK1-pY15/CDK1-total, respectively, sealed and stored at 4°C overnight. Anti-rabbit IgG Acceptor and anti-mouse IgG Donor beads were each added at a final concentration of 20 μg/ml and the reactions were incubated in the dark for 2 h at RT. Luminescence was measured using a Perkin Elmer EnVision Multimode plate reader with excitation at 680 nm and emission at 615 nm.

### Cell line-derived and patient-derived xenografts

HCC1569, OVCAR3 and SUM149PT cells were implanted at 5×10^6^ cells per mouse into the right flanks of female CB17-SCID, SCID-beige and NOD-SCID mice respectively (5-7 weeks old; Charles River), in 1:1 matrigel: media (Matrigel Corning, cat# CB35248). When tumors reached the target size of 100-150 mm^3^, (n=8) mice were randomized to treatment groups and treatment with RP-6306 was initiated. In vivo studies involving cell-derived xenografts were performed at Repare Therapeutics, in a CCAC (Canadian Council on Animal Care)-accredited vivarium with an Institutional Animal Care Committee-approved protocol.

In vivo studies using patient-derived xenografts (PDX) were conducted at Crown Biosciences Inc. (Taicang). Fresh primary human tumor tissue was harvested and cut into small pieces (approximately 2-3 mm in diameter). These tumor fragments were inoculated subcutaneously into the right flank of female BALB/c nude mice (5-7 weeks old) for tumor development and subsequently passaged by implantation into the cohort of mice enrolled in the efficacy study. Mice were randomized according to growth rate into treatment groups (n=6) when the mean tumor size reached approximately 150 (100–200) mm^3^. The procedures involving the care and use of animals in for the PDX model were reviewed and approved by the Institutional Animal Care and Use Committee (IACUC) of CrownBio prior to execution. During the study, the care and use of animals were conducted in accordance with the regulations of the Association for Assessment and Accreditation of Laboratory Animal Care (AAALAC).

RP-6306 was formulated in 0.5% methylcellulose and orally administered twice daily (BID, 0-8h) for a maximum of 21 days. Gemcitabine was administered once weekly intraperitoneally in saline. Animals were monitored for tumor volume, clinical signs and body weight three times per week. Tumor volume was measured using a digital caliper and calculated using the formula 0.52×L×W^2^. Response to treatment was evaluated for tumor growth inhibition (%TGI). TGI was defined as the formula: % TGI= ((TV_vehicle/last_ – TV_vehicle/day0_) - (TV_treated/last_ – TV_treated/day0_)) / (TV_vehicle/last_ – TV_vehicle/day0_) x100 calculated based on the means of the treatment groups at day 0 and last day of measurement. Change in body weight (BW) was calculated using the formula: % BW change = (BW_last_-BW_day0_/ BW_day0_) x100. BW change was calculated based on individual body weight changes relative to day 0. Statistical significance relative to vehicle control or other test groups was established by one-way ANOVA followed by Fisher’s LSD test for multiple groups and unpaired t-test for two group comparisons (GraphPad Prism v9.0).

### Blood and tumor tissue collection

Under isoflurane anesthesia, whole blood was collected by cardiac puncture and transferred to tubes containing 0.1 M citric acid (3:1 citric acid:blood) and stored at −20°C for LC-MS/MS analysis.. Tumors were removed from mice flanks and cleared of surrounding mouse stroma. Tumor pieces between 50 mg and 100 mg were collected in a pre-weighed pre-filled bead mill tube (Fisher Scientific, Cat# 15-340-154) and then flash-frozen in liquid nitrogen. Other tumor fragments from vehicle- and compound-treated mice were placed in 10% neutral buffered formalin (NBF) within 2-5 minutes of surgical excision, fixed in NBF for 18-24 hours at room temperature and embedded in paraffin.

### RP-6306 Quantitation by LC-MS-MS

The extraction of whole blood samples was performed by protein precipitation using four volumes of acetonitrile. The sample extracts were analyzed using a Transcend LX2 / Ultimate 3000 liquid chromatography system coupled to a Thermo Altis triple quadrupole electrospray mass spectrometer (Thermo Fisher Scientific) operated in positive mode. Separations were performed using a 2 x 50mm, 2.8µm Pursuit XRS C8 HPLC column (Agilent). A reversed-phase linear gradient of water + 0.1% formic acid and 1:1 acetonitrile:MeOH was used to elute RP-6306 and the internal standard. Samples were quantified against a 10-point linear standard curve and 3 levels of quality control samples. Whole blood concentrations of RP-6306 were converted to free unbound plasma concentrations using an in vitro derived blood / plasma ratio = 1.2 and fraction unbound (f_u_) plasma = 0.185 from the CD-1 mouse strain.

### Immunohistochemistry

Histology in Fig. S10 was performed by HistoWiz Inc. Briefly, the formalin-fixed tissues were dehydrated through a 20%, 80%, 95% and 100 % ethanol series, cleaned in xylene, embedded in paraffin then sectioned at 4 μm. Immunohistochemistry for *γ*H2AX, CDK1pT14 and cyclin B1-pS126 were performed on a Bond Rx autostainer (Leica Biosystems) with heat antigen retrieval. Bond polymer refine detection (Leica Biosystems) was used according to manufacturer’s protocol. After staining, sections were dehydrated and film coverslipped using a TissueTek-Prisma and Coverslipper (Sakura). Whole slide scanning (40x) was performed on an Aperio AT2 (Leica Biosystems). Image quantification analysis was performed using HALO. H-score is given by the formula: H-score = (1x % weak positive cells) + (2x % moderate positive cells) + (3x % strong positive cells). Histology in Fig. S11c was performed by NeoGenomics Inc. Briefly, formalin fixed, paraffin embedded tumor samples were sectioned at 4 μm, mounted on charged glass slides and baked at 60°C for 1 h. Immunohistochemistry for cyclin E1 was performed on a Bond-III autostainer (Leica Biosystems). Bond polymer refine detection (Leica Biosystems) was used according to manufacturer’s protocol. Slides were then removed from the instrument dehydrated, cleared, coverslipped. Brightfield images (20x) were acquired on an Aperio AT2 (Leica Biosystems).

### ELISA assay

Tumor samples were homogenized in MSD Tris lysis buffer (Meso Scale Discovery, #R60TX-2) supplemented with 1X Halt Protease (Thermo Fisher Scientific, #78429) and phosphatase inhibitors (Thermo Fisher Scientific, #78426) using a Beadruptor tissue homogenizer (OMNI International. After homogenization, samples were centrifuged at 14000x *g* for 5 min at 4°C. ELISA plates were coated with the capture antibody (CDK1) incubated overnight at 4°C, washed and then blocked for 1 h at room temperature. Tissue samples (60 μg) were added to the plates to incubate for 2.5 h at room temperature. After washing, the detector antibody (CDK1-pT14) was added for 1 h at room temperature. After washing and plate drying, detection occurred using a secondary anti-rabbit HRP conjugate incubation for 1 h followed by a 10-min incubation with TMB peroxidase substrate stop solution (Thermo Fisher Scientific, #N600). The absorbance was measured in 96-well plate format on an EnVision2105 at 450 nm. Samples were quantified relative to a standard protein extract and an MSD lysis buffer used as a blank to control for inter-day variability.

## Acknowledgments

We are grateful to Frank Sicheri, Agnel Sfeir and Marcel van Vugt for critical reading of the manuscript. We also thank Jiri Lukas, Philipp Kaldis and David Pellman for helpful comments and for Crown Bio for allowing us to report copy number and genomic information for the PA1443 tumor. We thank Jason Moffat for his generous sharing of the TKO sgRNA libraries, Ronny Drapkin for the FT282 cells, Ralph Scully for the pHIV-NAT plasmid, Monica Hasegan and Kin Chan at the NBCC (LTRI) for high content imaging support and Illumina sequencing respectively. We also thank Charles Pellerin for help with assay development. AAQ is a recipient of a CIHR postdoctoral fellowship, DD is a Canada Research Chair (Tier I) and work in the DD lab was supported by grants from the CIHR (FDN143343) and Repare Therapeutics.

---

**Supplementary figure 1. Related to figure 1.**
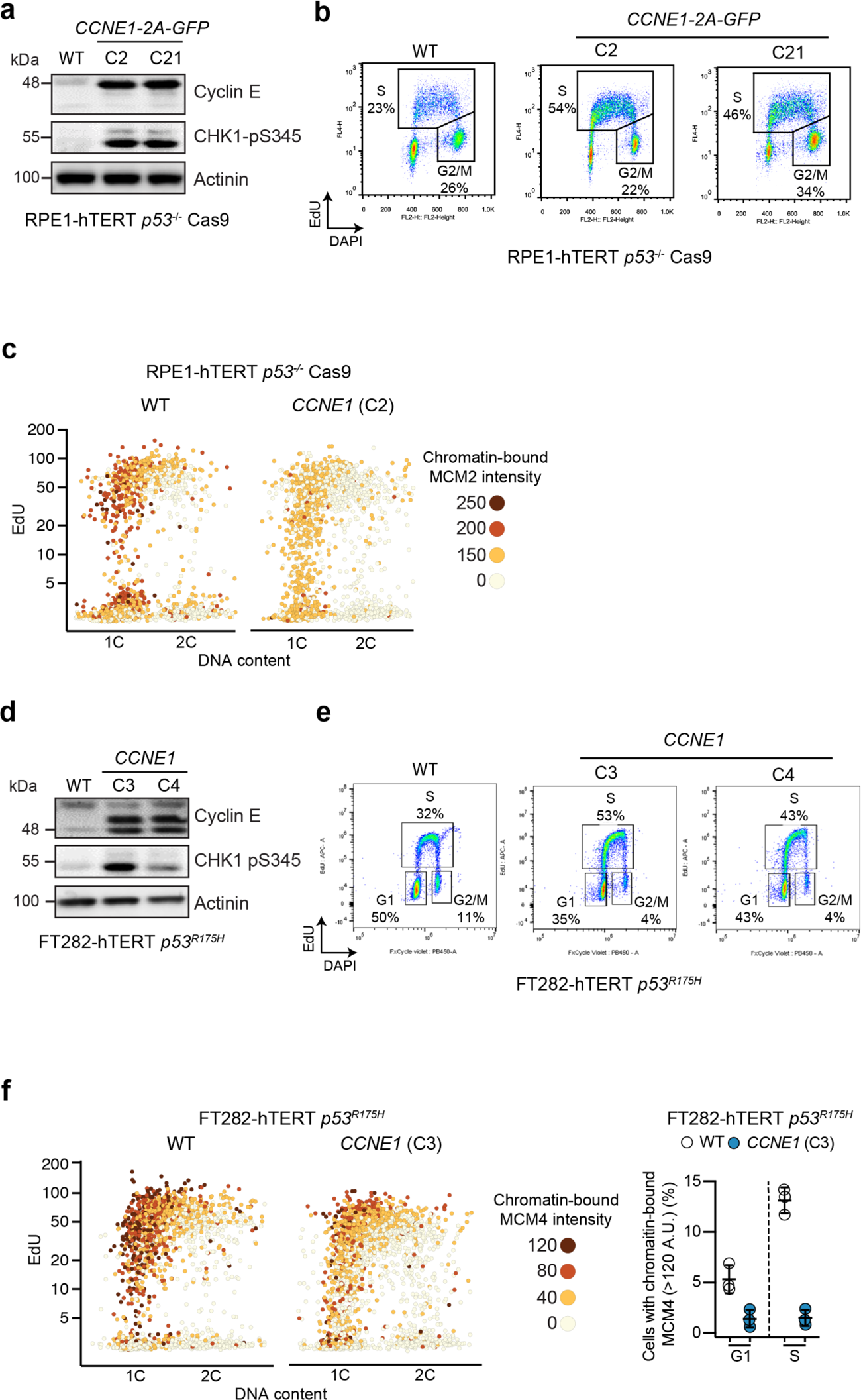
**a,d**, Whole cell lysates of (a) RPE1-hTERT *p53^-/-^* Cas9 and (d) FT282-hTERT *p53^R175H^ CCNE1-*high and parental (WT) cells were immunoblotted with cyclin E, CHK1-pS345 and actinin specific antibodies. **b,** FACS analysis of RPE1-hTERT *p53^-/-^* Cas9 *CCNE1-*high and parental (WT) cells for EdU incorporation and DNA content (DAPI). Percentage of each population gated for EdU^+^ (S) and EdU-2C (G2/M) are indicated. **c.** QIBC analysis of RPE1-hTERT *p53^-/-^* Cas9 *CCNE1-*high and parental cells for chromatin-bound MCM2 nuclear intensity, EdU incorporation and DNA content (DAPI). **e,** FACS analysis FT282-hTERT *p53^R175H^ CCNE1-*high and parental (WT) cells for EdU incorporation and DNA content (DAPI). Percentage of each population gated for EdU^-^ 1C (G1), EdU^+^ (S) and EdU-2C (G2/M) are indicated. **f,** Left, QIBC analysis of FT282-hTERT *p53^R175H^ CCNE1-*high and parental cells for chromatin-bound MCM4 nuclear intensity, EdU incorporation and DNA content (DAPI). Right, quantitation of EdU^-^ 1C (G1) and EdU^+^ (S) nuclei with chromatin-bound MCM4 (A.U. > 120). Data are shown as mean ± S.D. (n=3).

**Supplementary figure 2. Related to figure 1.**
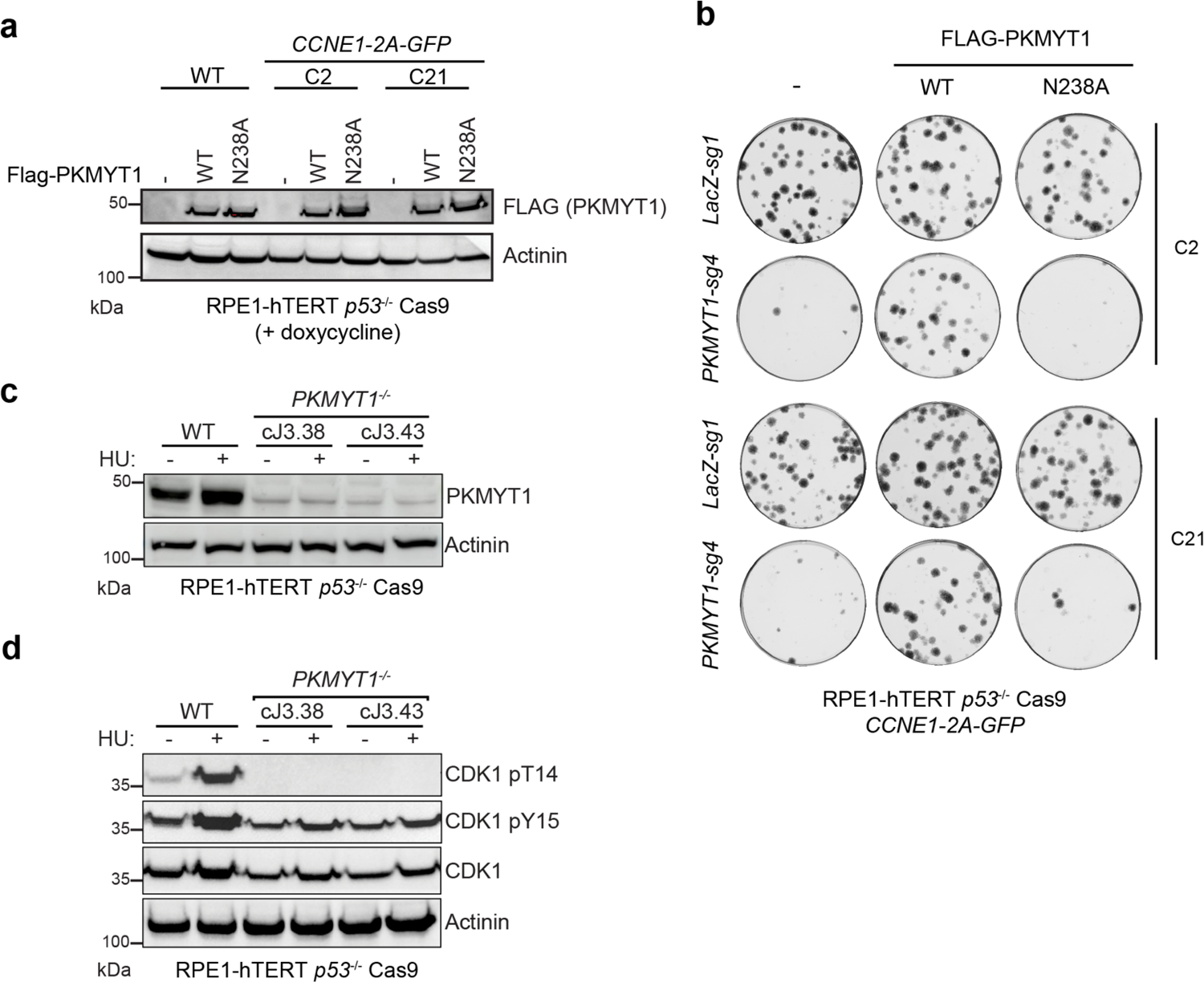
**a**, Whole cell lysates of RPE1-hTERT *p53^-/-^* Cas9 *CCNE1-*high and parental cells expressing doxycycline-inducible sgRNA-resistant FLAG alone (-), *FLAG-PKMYT1* or *FLAG-PKMYT1-N238A* were immunoblotted with FLAG and actinin antibodies. **b,** Representative plates stained with crystal violet from clonogenic survival assays with *CCNE1-high* cells transduced with lentivirus expressing the indicated sgRNAs. **c,d,** Whole cell lysates of RPE1-hTERT *p53^-/-^* Cas9 parental and two independent *PKMYT1^-/-^* clones either untreated or treated for 24 h with 3 mM hydroxyurea (HU) were immunoblotted for PKMYT1 (c) and either total CDK1, CDK1-pT14 or CDK-pY15 (d). Actinin was used as loading control.

**Supplementary figure 3. Related to figure 2.**
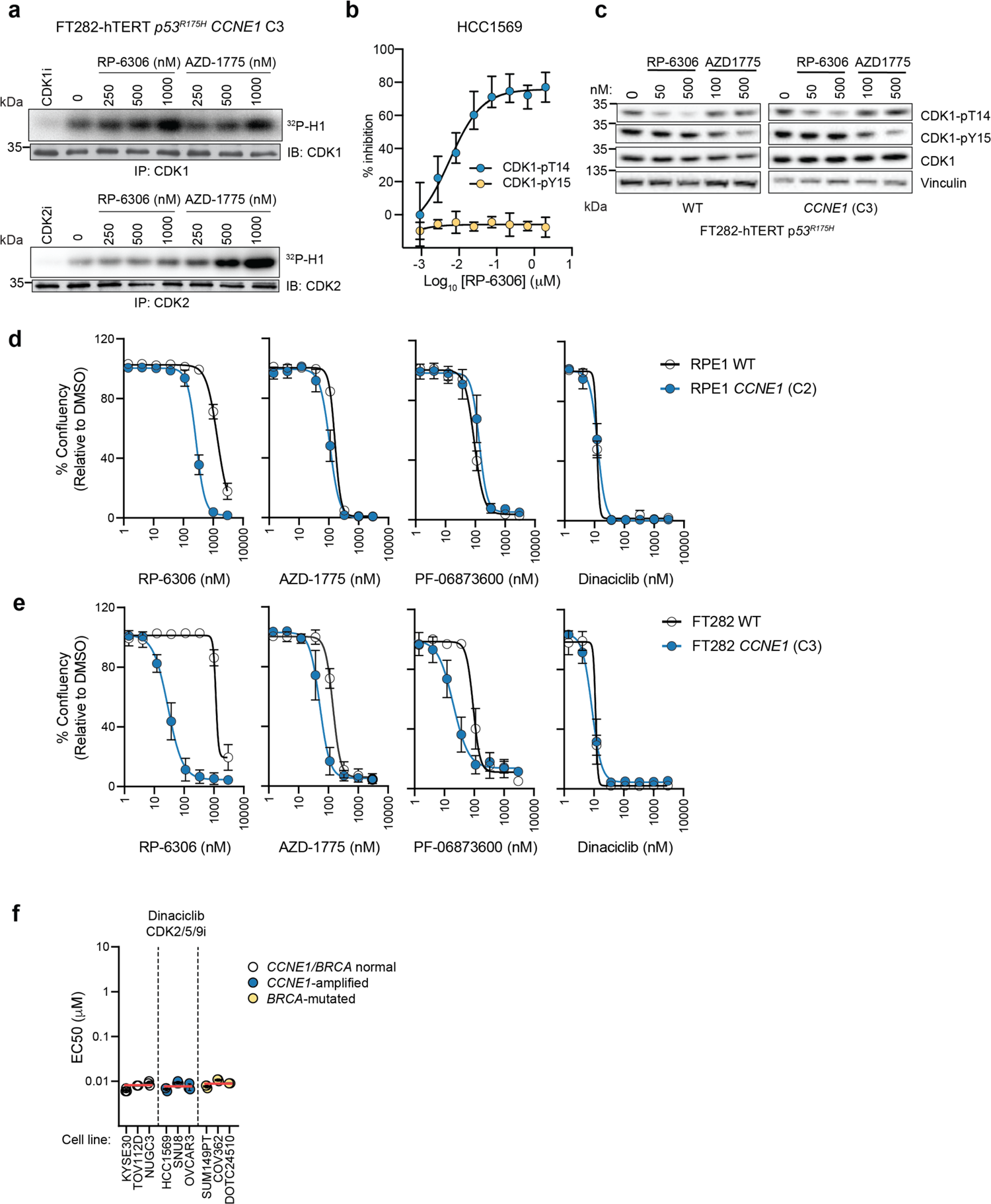
**a**, Cell culture of FT282-hTERT *p53^R175H^ CCNE1-high* cells were treated with the indicated doses of RP-6306 and AZD-1775 for 24 h. Cellular extracts were prepared and immunoprecipitated (IP) with agarose-coupled CDK1 (upper) or CDK2 (lower) antibodies. Immunoprecipitates were subjected to in vitro kinase assays using [*γ*^32^P]-ATP and recombinant histone H1 as a substrate. Reactions were resolved by SDS-PAGE and gels were imaged using a phosphor screen. A sample of each immunoprecipitate was immunoblotted (IB) with CDK1 or CDK2 antibodies. CDK1i (RO-3306) or CDK2i (PF-06873600) was added to indicated in vitro reactions. **b.** HCC1569 cells were treated with the indicated doses of RP-6306 for 2 h and subjected to the AlphaLISA assay using CDK1-pT14, CDK1-pY15 and total CDK1 antibodies. Data are shown as mean ± S.D. (n=3). **c,** FT282-hTERT *p53^R175H^* parental and *CCNE1-*high cells were treated with the indicated doses of RP-6306 and AZD-1775 for 4 h. Whole cell lysates were prepared and immunoblotted with antibodies against CDK1-pT14, CDK1-pY15, CDK1 and vinculin as a loading contol. **d,e,** EC50 determination for growth inhibition for the parental and *CCNE1*-high cells in the (d) RPE1-hTERT *p53 ^-/-^* Cas9 (RPE1) (e) FT282 *p53^R175H^* (FT282) and (g) indicated cancer cell lines treated with the indicated compounds. Growth was monitored with an Incucyte for up to 6 population doublings. Data are shown as mean ± S.D. (n=3). In (f) cell lines are also grouped according to their *CCNE1* or *BRCA* status and the red bar indicates the mean of each grouping.

**Supplementary figure 4. Related to figure 3.**
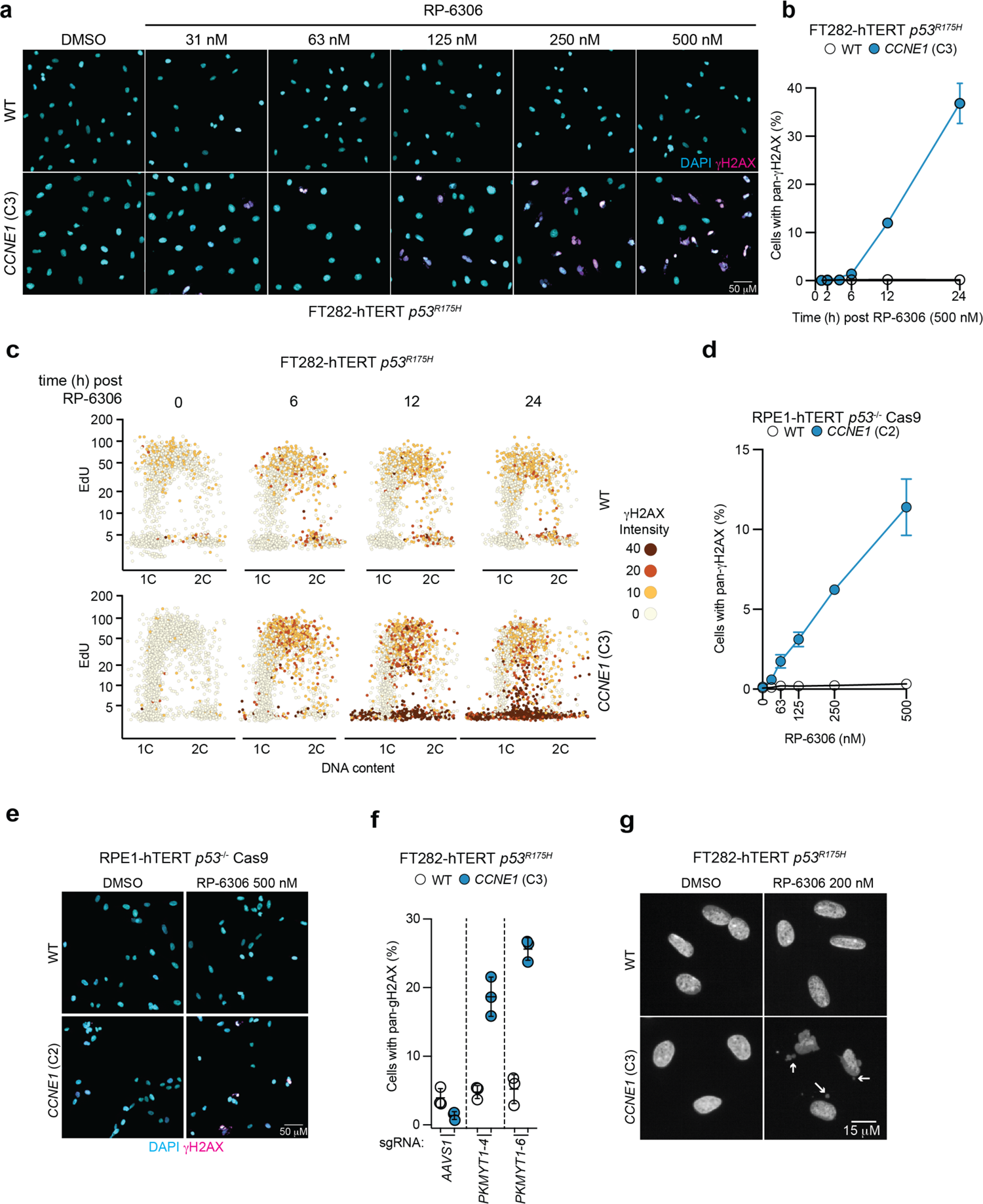
**a**, Representative QIBC micrographs of FT282-hTERT *p53^R175H^* (WT) and *CCNE1-*overexpressing (*CCNE1*) cells treated either with DMSO or increasing doses of RP-6306. The DAPI (cyan) and *γ*H2AX (magenta) channels are merged. **b,c,** QIBC analysis of *γ*H2AX nuclear intensity, EdU incorporation and DNA content (DAPI) with the quantitation of cells with pan-*γ*H2AX as a function of time after addition of RP-6306 (500 nM) shown in (b) and representative QIBC plots shown in (c). Data are shown as mean ± S.D. (n=3). **d,e,** QIBC analysis of *γ*H2AX nuclear intensity of RPE1-hTERT *p53^-/-^* Cas9 parental (WT) and *CCNE1-2A-GFP* (*CCNE1*) cells. Quantitation of cells with pan-*γ*H2AX as a function of RP-6306 dose is shown in (d) and representative micrographs of cells treated with DMSO or 500 nM RP-6306 with the DAPI (cyan) and *γ*H2AX (magenta) channels merged shown in (e). Data are shown as mean ± S.D. (n=3). **f,** QIBC Quantitation of cells with pan-*γ*H2AX after transduction with lentivirus expressing the indicated sgRNAs. **g**, Representative micrographs of cells with micronuclei (white arrows) in FT282 parental and *CCNE1-*high cells following treatment with either DMSO or RP-6306 (200 nM) for 72 h.

**Supplementary figure 5. Related to figure 3.**
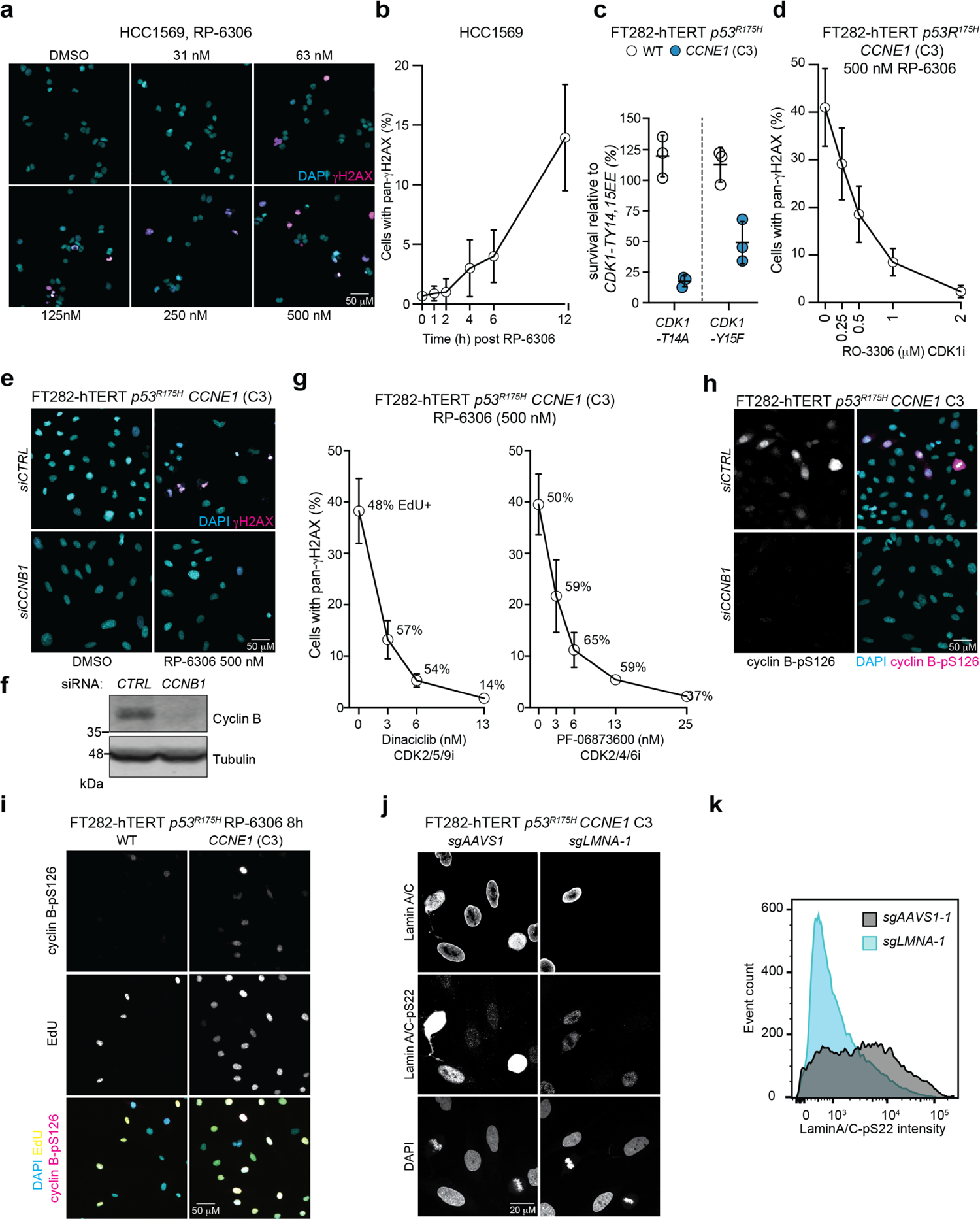
**a**, Representative QIBC micrographs of HCC1569 cells treated with increasing doses of RP-6306. The DAPI (cyan) and *γ*H2AX (magenta) channels are merged. **b,** QIBC quantitation of pan-*γ*H2AX staining as a function of time after addition of RP-6306 (500 nM) in HCC1569 cells. **c,** Clonogenic survival assays of the indicated FT282-hTERT *p53^R175H^* Cas9 cell lines transduced with lentivirus expressing CDK1-T14A-GFP or CDK1-Y15F-GFP relative to CDK1-TY14,15EE-GFP. Data are shown as mean ± S.D. (n=3). **d,** QIBC quantitation of pan-*γ*H2AX staining in FT282-hTERT *p53^R175H^ CCNE1* cells treated with RP-6306 (500 nM) as a function of CDK1 inhibitor RO-3306 dose. Data are shown as mean ± S.D. (n=3). **e,f,** FT282-hTERT *p53^R175H^ CCNE1* cells transfected with either non-targeting siRNA (*siCTRL*) or siRNA targeting cyclin B (*siCCNB1)* were treated with RP-6306 (500 nM). Representative QIBC micrographs with DAPI (cyan) and *γ*H2AX (magenta) channels merged are shown in (e). Immunoblot analysis of cyclin B levels in lysates prepared from DMSO-treated cells is shown in (f). Tubulin was used as a loading control. **g,** QIBC quantitation of pan-*γ*H2AX staining in FT282-hTERT *p53^R175H^ CCNE1* treated with RP-6306 (500 nM) as a function of the dose of dinaciclib or (left) PF-06873600 (right). Data are shown as mean ± S.D. (n=3). **h**, Representative QIBC micrographs of FT282-hTERT *p53^R175H^* (WT) and *CCNE1-*overexpressing (*CCNE1*) cells transfected with the indicated siRNA and stained with DAPI (cyan) and cyclin B-pS126 antibody (magenta). The channels are merged. **i,** Representative QIBC micrographs of FT282-hTERT *p53^R175H^* (WT) and *CCNE1-*overexpressing (*CCNE1*) cells treated with RP-6306 (500 nM) and stained with DAPI (cyan), EdU (yellow) and a cyclin B-pS126 antibody, (magenta). The channels are merged. **j,k,** Validation of the lamin A/C-pS22 antibody. Representative micrographs of FT282-hTERT *p53^R175H^ CCNE1* cells transduced with lentivirus expressing sgRNAs targeting either *AAVS1* (*sgAAVS1*) or *LMNA* (*sgLMNA-1*) stained with DAPI and the indicated antibodies (j). Flow cytometry histograms of the same cells showing loss of Lamin A/C-pS22 signal in the *sgLMNA-1* condition.

**Supplementary figure 6. Related to figure 3.**
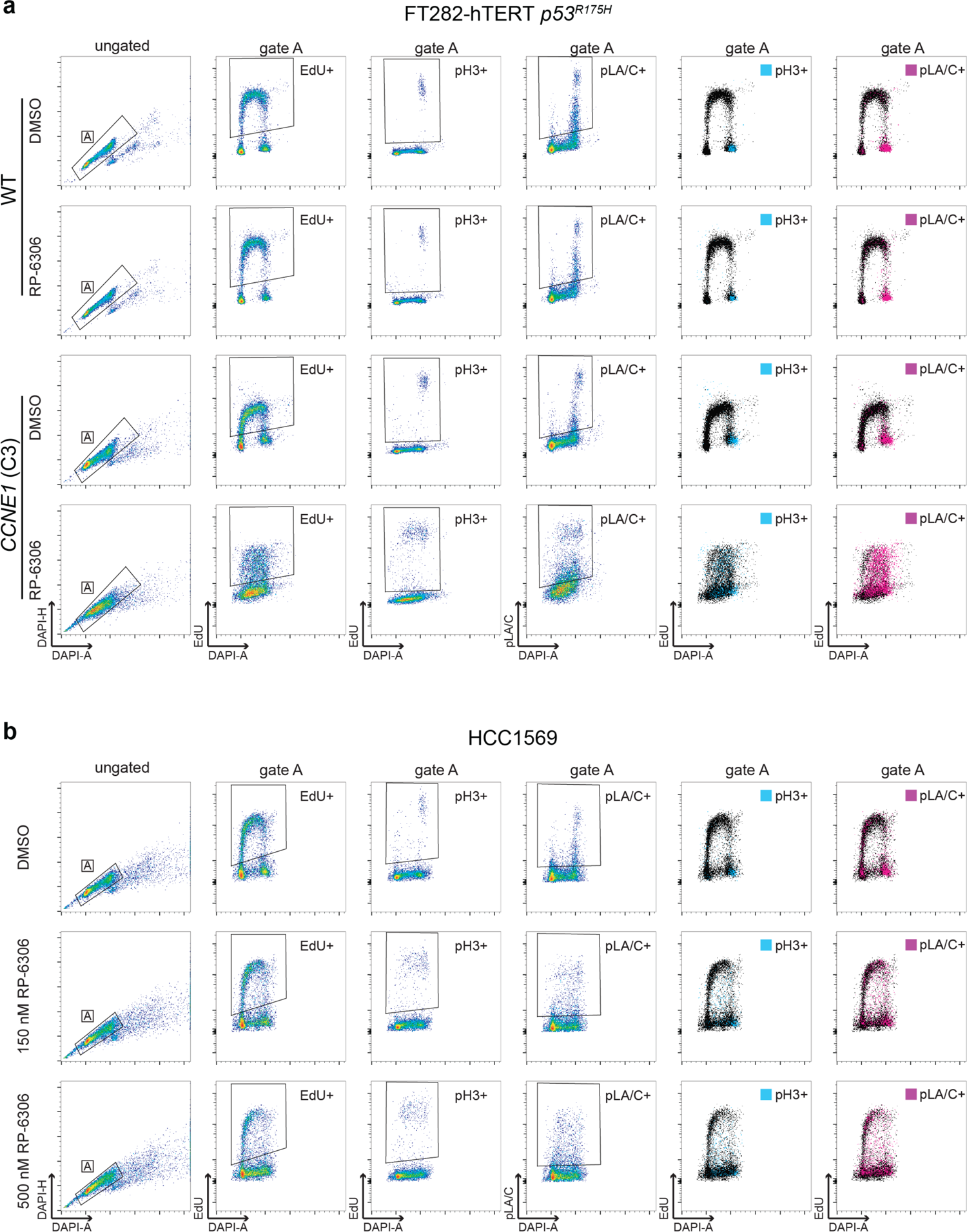
**a,b,**, Gating strategy for quantitation of EdU^+^ H3-pS10^+^ (pH3^+^) and lamin A/C-pS22^+^ (pLA/C^+^) FT282-hTERT *p53^R175H^* (a) and HCC1569 (b) cells by FACS. Single cells were identified by gating events on DAPI-H/DAPI-A (gate A). EdU/DAPI, pH3/DAPI and pLA/C/DAPI were then plotted and EdU^+^, H3-pS10^+^ and lamin A/C-pS22^+^ populations were identified relative to unlabeled controls.

**Supplementary figure 7. Related to figure 4.**
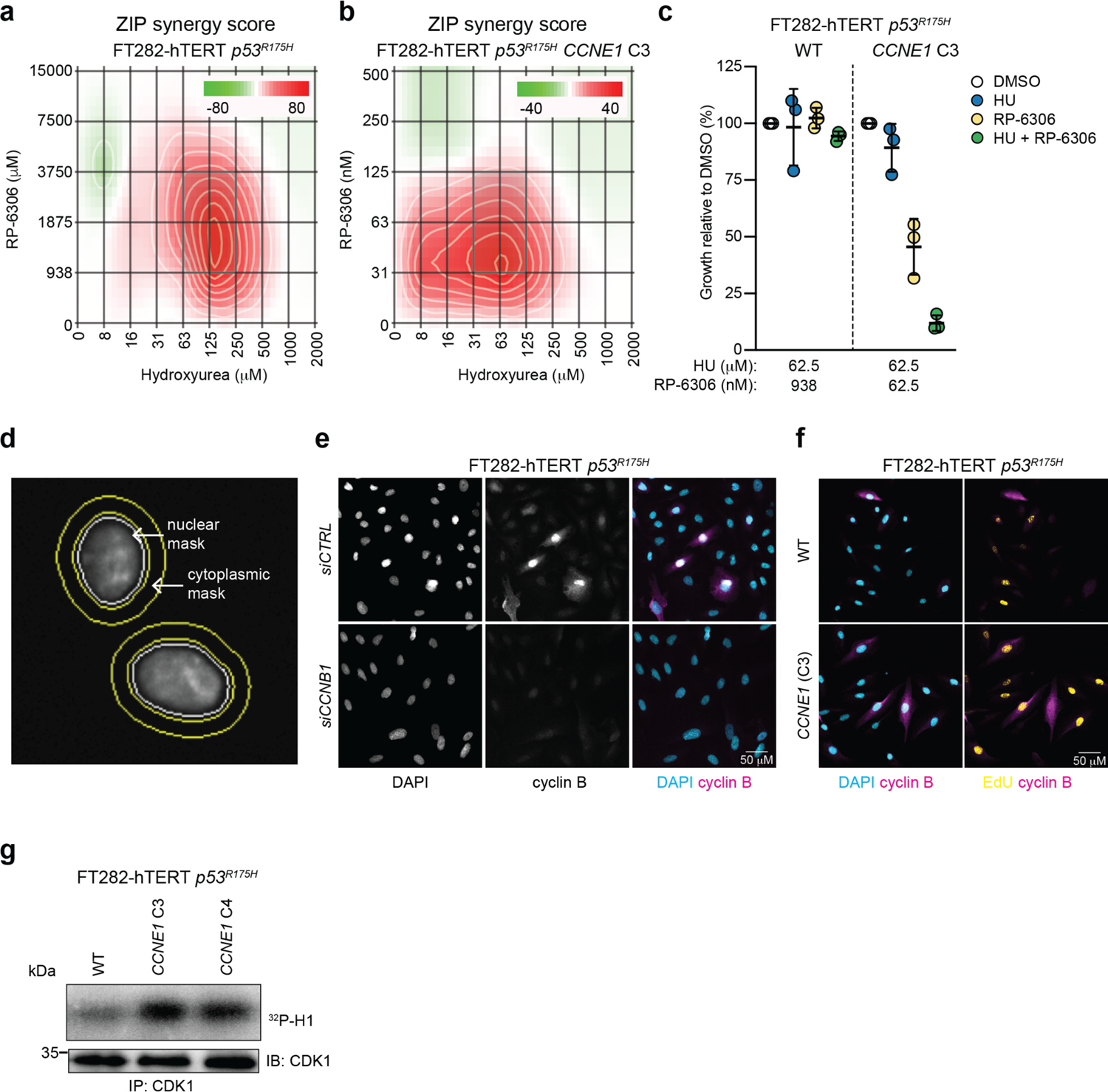
**a,b**, ZIP synergy scores at various dose combinations of RP-6306 and hydroxyurea (HU) in FT282-hTERT *p53^R175H^* parental (WT) (a) and *CCNE1-*high (b) cells. Score ≥10 (red color) represents synergy, ≤-10 (green) represents antagonism. Values were obtained by analyzing mean data from 3 independent biological replicates with SynergyFinder. Growth was monitored with an Incucyte for up to 6 population doublings. **c,** Growth inhibition relative to DMSO control of parental (WT) and *CCNE1-*high cells after treatment with the indicated dose of hydroxyurea (HU), RP-6306 or the combination of both. Growth was monitored with an Incucyte for up to 6 population doublings. Data are shown as mean ± S.D. (n=3). **d,** Approach for cytoplasmic cyclin B quantitation by QIBC. The nuclear mask edge was expanded by 8 and 28 pixels to create a doughnut mask used to estimate cytoplasmic signal intensity. **e,** Representative micrographs of FT282-hTERT *p53^R175H^ CCNE1-*high cells transfected with the indicated siRNAs and stained with DAPI (cyan) and cyclin B antibodies (magenta). The channels are merged. **f,** Representative QIBC micrographs of FT282-hTERT *p53^R175H^* parental (WT) and *CCNE1-*high cells stained with cyclin B antibodies (magenta) and either DAPI (cyan) or EdU (yellow) channels merged. **f,** Cellular extracts of the indicated cell lines were prepared and immunoprecipitated with agarose-coupled CDK1 antibodies. Immunoprecipitates (IP) were subjected to in vitro kinases assays using [*γ*-^32^P]-ATP and recombinant histone H1 as a substrate. Reactions were resolved by SDS-PAGE and gels were imaged using a phosphor screen. A sample of each immunoprecipate was immunoblotted (IB) with a CDK1 antibody as loading control.

**Supplementary figure 8. Related to figure 4.**
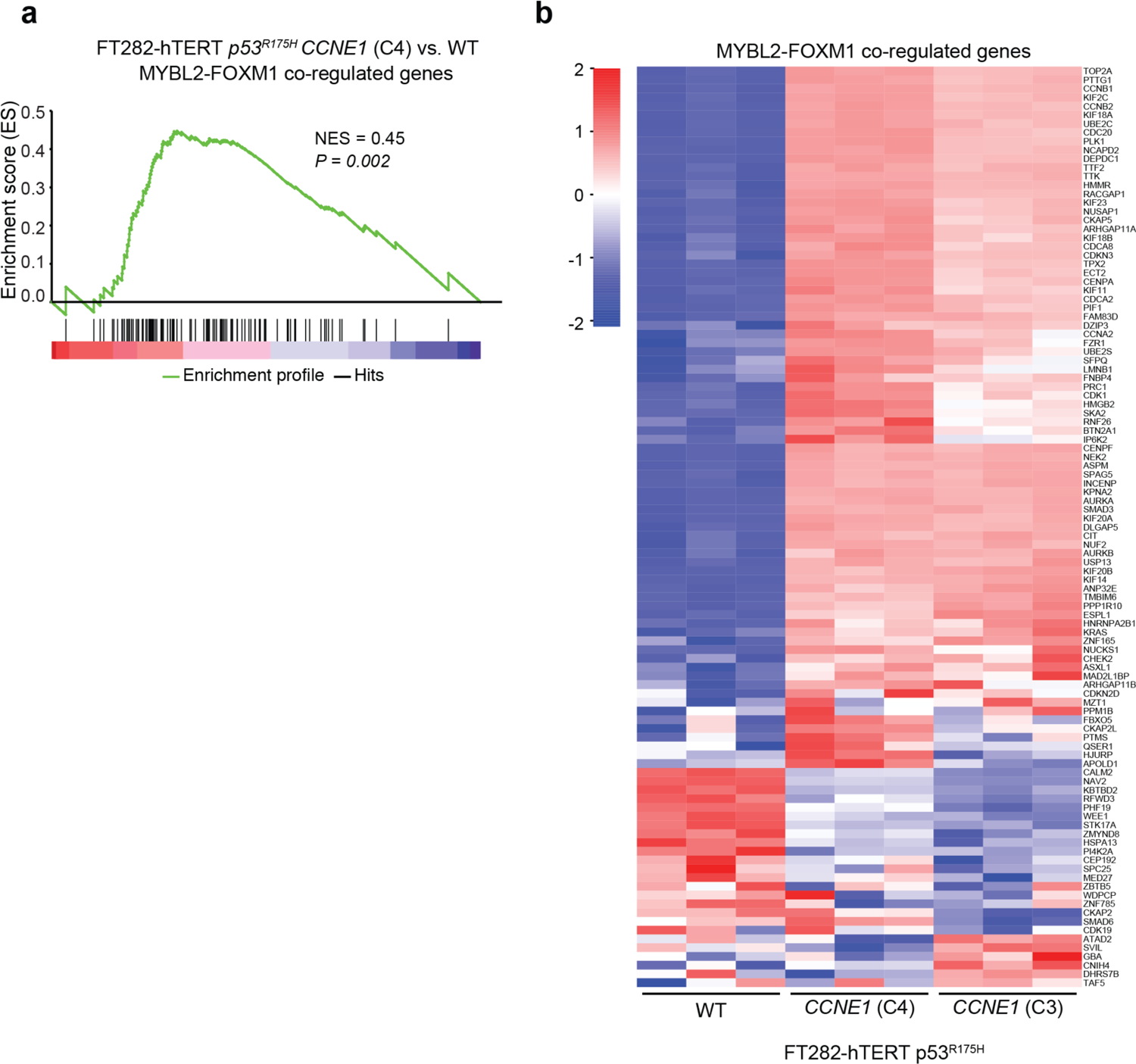
**a,b**, Gene set enrichment analysis (GSEA) (a) and heat map (b) of differential gene expression in FT282 parental (WT) vs *CCNE1*-high (C4) cells for a gene set comprising genes co-regulated by MYBL2 and FOXM1 (Table S5). The heat map also includes FT282 parental vs *CCNE1*-high (C3).

**Supplementary figure 9. Related to figure 5.**
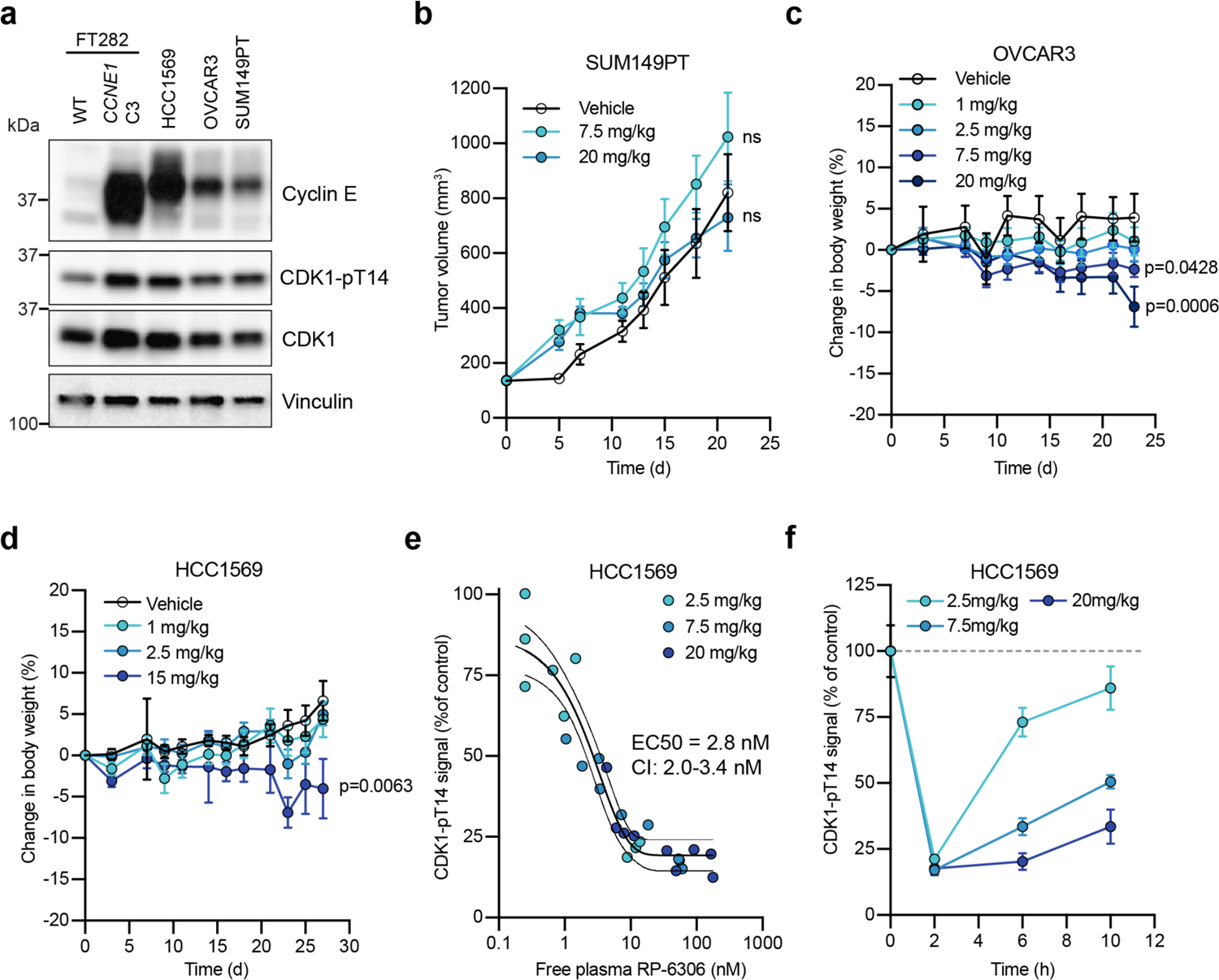
**a**, Whole cell lysates of the indicated cell lines were immunoblotted with antibodies against cyclin E, CDK1-pT14, CDK1 and vinculin. **b-d**, RP-6306 was administered orally BID at the indicated doses for the duration of the experiment. **b,** Tumor growth of SUM149PT xenografts in NOD-SCID mice treated with either RP-6306 or vehicle BID for the duration of the experiment. Results are expressed as mean tumor volume ± SEM (n=8). n.s. = non-significant p-value **c, d**, Changes in body weight in tumor-bearing OVCAR3 (c) and HCC1569 (d) CB-17 SCID and SCID-beige mice treated with either RP-6306 or vehicle. Results are expressed as mean % body weight change ± SEM (n=8). Significant p-values are indicated. **e, f,** Mice bearing HCC1569 tumors were treated with RP-6306 at the indicated doses BID for 1.5 days and tumor tissue and whole blood sampled at 2, 6 and 10 h post last dose. Data represent the free (unbound to plasma protein) blood concentration of RP-6306 for mouse relative to tumor CDK1-pT14 signal inhibition quantified by ELISA relative to vehicle-treated control tumors (N=3/group/time point). **(e)** The tumor EC50 was determined by a non-linear least square’s regression to a normalized dose-response model with 95% confidence intervals. **f,** Kinetics of CDK1-pThr14 inhibition with time. Data presented as the mean ± SEM (n=3).

**Supplementary figure 10. Related to figure 5.**
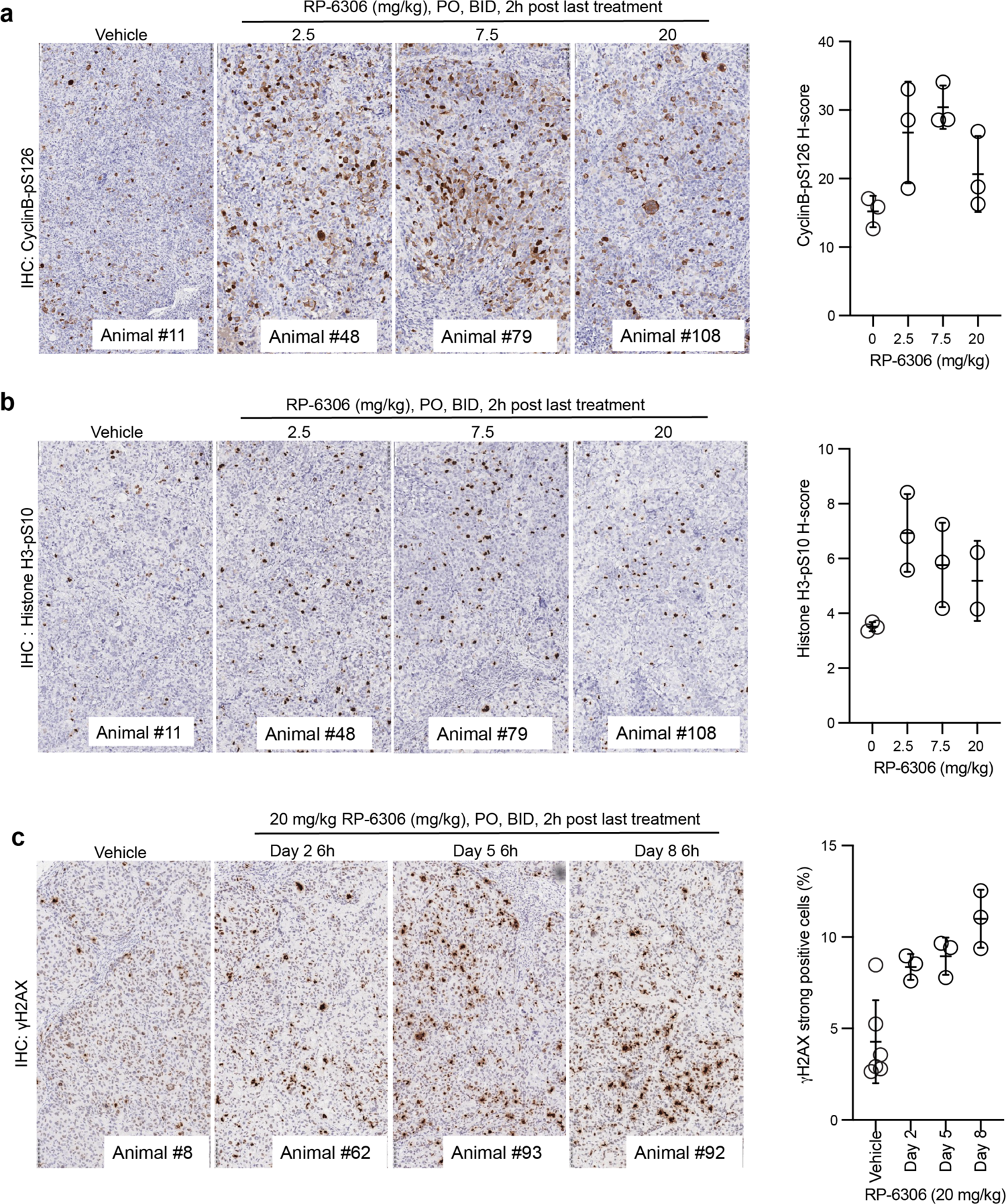
**a,b**, HCC1569 tumor-bearing mice were administered RP-6306 orally BID for 1.5 days and sacrificed at 2 h post last treatment and tumor tissue prepared for FFPE. Tumor tissues were stained with cyclin B-pS126 (a) or histone H3-pS10 (b) and the percentage of positive staining was quantified by HALO software. Results are expressed as mean H-score ± SEM (n=3). **c**, HCC1569 tumor-bearing mice were treated with 20 mg/kg RP-6306 or vehicle BID and were sacrificed 6 h post treatment at the indicated times and prepared for FFPE. Tumor tissues were stained with *γ*H2AX and the percentage of strong positive *γ*H2AX cells present in the tumor area was quantified by HALO software. Results are expressed as mean ± SEM (n=3).

**Supplementary figure 11. Related to figure 5.**
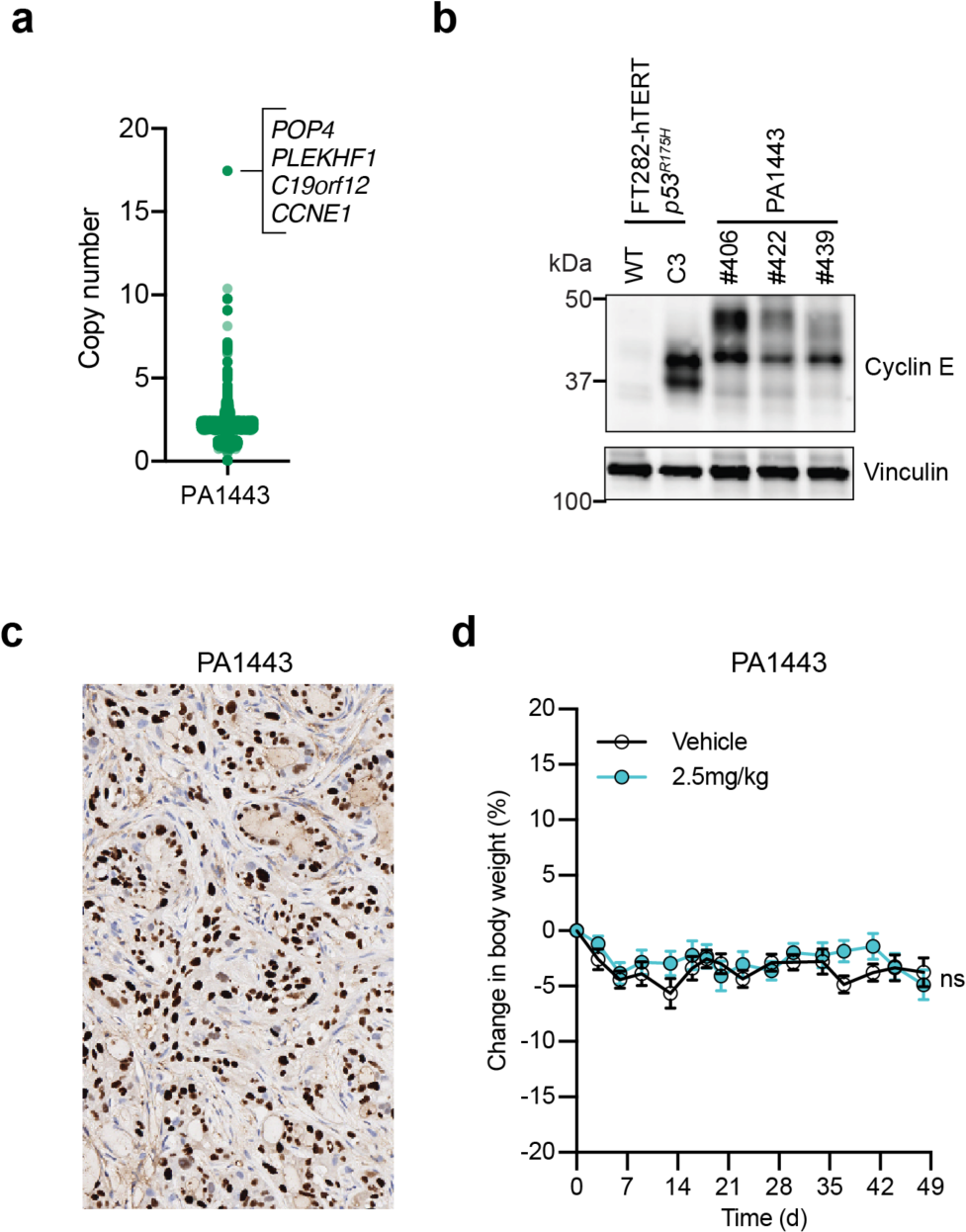
**a**, Distribution of the gene-level copy number in the *CCNE1*-amplified pancreatic cancer (PA1443) patient-derived xenograft (PDX). Highlighted is the amplicon containing *CCNE1*. **b,** Whole cell lysates from FT282-hTERT *p53^R175H^* parental (WT) and *CCNE1*-high (C3) cell lines and PA1443 PDX tumor tissue were immunoblotted with antibodies to cyclin E and vinculin. **c,** Tumor tissues of PA1443 PDX implanted in BALB/c nude mice were prepared for FFPE and stained with a cyclin E1 antibody. **d,** Changes in body weight in tumor-bearing PA1443 PDX implanted in BALB/c nude mice treated either with RP-6306 or vehicle. RP-6306 was administered orally BID at 2.5 mg/kg for the duration of the experiment. Results are expressed as mean % body weight ± SEM (n=8). *p* value relative to vehicle was determined with a one-way ANOVA test. Only p values <0.05 are indicated. ns, not significant (*p* value > 0.05)

**Supplementary figure 12. Related to figure 5.**
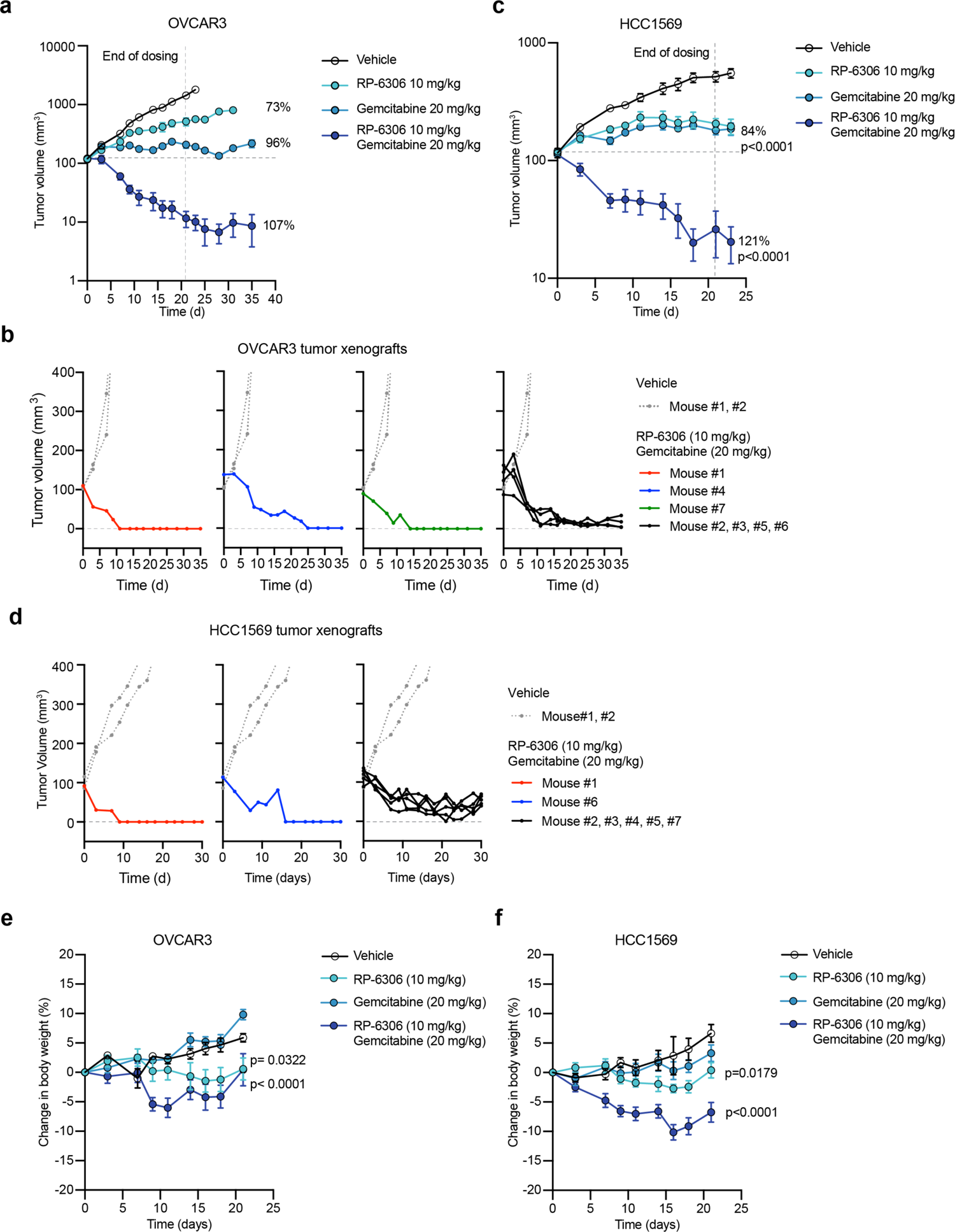
**a, c**, Same data as presented in Fig 5d,e with tumour growth values plotted in log scale for OVCAR3 (a) and HCC1569 (c). Please refer to Fig 5 for TGI and *p* values. **b, d,** Growth traces for individual OVCAR3 (b) and HCC1569 (d) tumors from the experiments shown in Fig. 5d,e from mice treated with the gemcitabine/RP-6306 combination. For comparison, tumor growth data for two mice treated with vehicle are shown. **e,f,** Changes in body weight in tumor-bearing OVCAR3 (e) CB-17 SCID and HCC1569 SCID-beige (f) mice treated with either RP-6306, gemcitabine or both. Gemcitabine was delivered once weekly intraperitoneally and RP-6306 was given orally BID for 21 d after which all treatments were stopped, and body weight monitored for the remainder of the experiment. Results are expressed as mean ± SEM (n=8). Also indicated are *p* values relative to vehicle were determined with a one-way ANOVA test. Only *p* values <0.05 are indicated.

**Supplementary figure 13.**
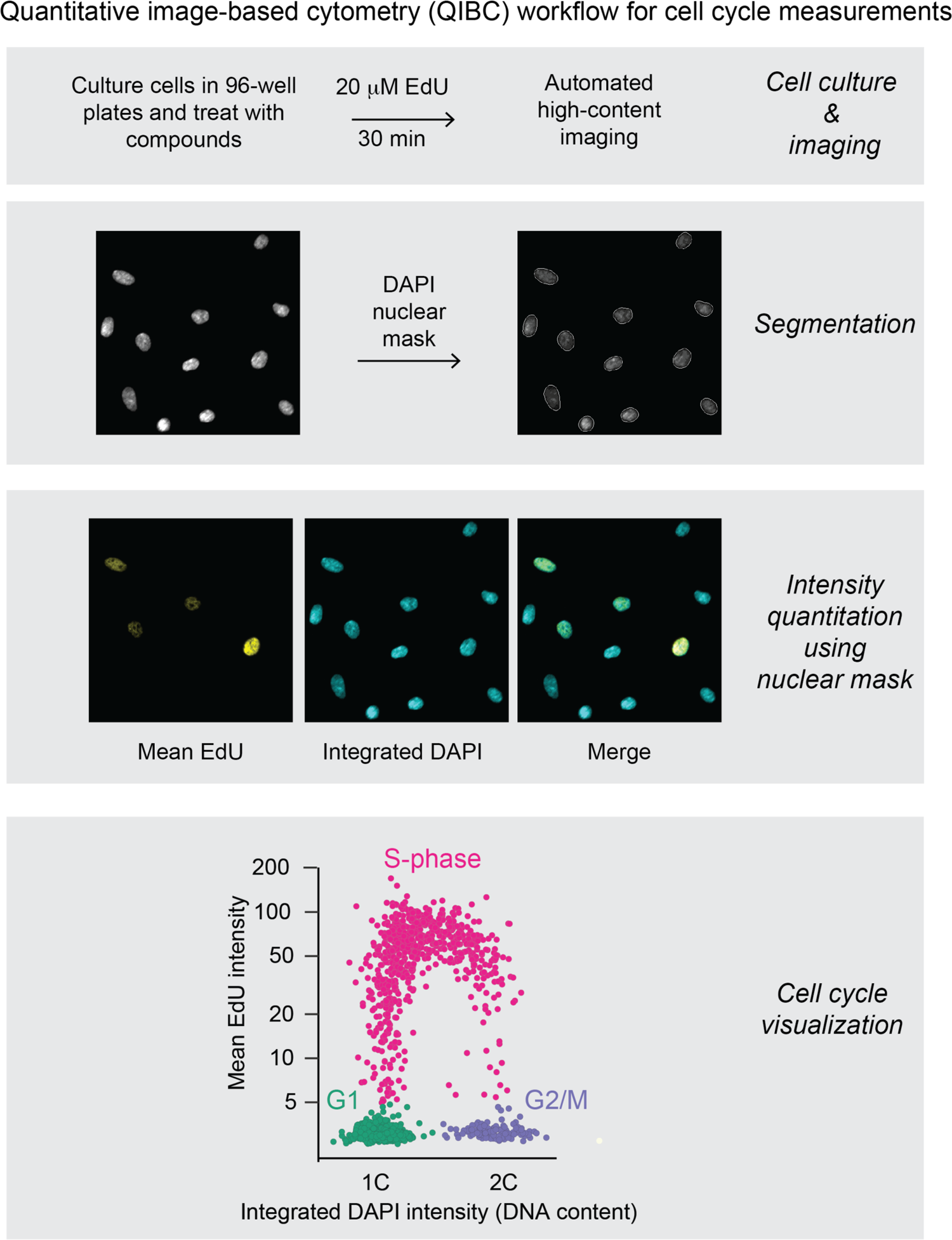
Quantitative image-based cytometry (QIBC) workflow. Cells were plated in 96-well plates and subjected to desired experimental treatments. Before harvesting, cells were pulsed with 20 μM EdU for 30 min to label nascent DNA. Plates were subjected to immunofluorescence analysis with antibodies directed against proteins and post-translational modifications of interest, DAPI to stain DNA and click chemistry to detect EdU incorporation in cells undergoing DNA replication. Following high-content image acquisition, nuclei were segmented using the DAPI channel and a nuclear mask was applied to each channel to quantitate the integrated DAPI intensity, mean EdU intensity and mean intensity of staining with other antibodies. The low and high end of the DAPI distribution corresponds to one genome copy (1C) or two genome copies (2C) respectively. To visualize cell cycle distributions, mean EdU intensity was plotted as a function of integrated DAPI intensity which allowed for clear identification of cells in G1 (1C, EdU^-^), S-phase (EdU^+^) and G2/M (2C, EdU^-^).

## Supplementary Tables

**Supplementary Tables 1 & 2** will be available at publication

**Supplementary Table 3.** Summary of TIDE editing analyses performed in the course of this study

| Experiment | Cell line | Genotype | sgRNA used | indel score (%) |
| --- | --- | --- | --- | --- |
| Related to Fig. 1d-e | RPE1 hTERT Cas9 *p53*<sup>-/-</sup> | WT | *PKMYT1-4* | 85 |
|  |  |  | *PKMYT1-6* | 91 |
|  |  |  | *WEE1-1* | 94 |
|  |  |  | *WEE1-3* | 91 |
|  |  |  | *CDK2-1* | 81 |
|  |  |  | *CDK2-2* | 96 |
|  |  | *CCNE1-2A-GFP* (C2) | *PKMYT1-4* | 89 |
|  |  |  | *PKMYT1-6* | 93 |
|  |  |  | *WEE1-1* | 91 |
|  |  |  | *WEE1-3* | 86 |
|  |  |  | *CDK2-1* | 77 |
|  |  |  | *CDK2-2* | 93 |
|  |  | *CCNE1-2A-GFP* (C21) | *PKMYT1-4* | 81 |
|  |  |  | *PKMYT1-6* | 97 |
|  |  |  | *WEE1-1* | 89 |
|  |  |  | *WEE1-3* | 96 |
|  |  |  | *CDK2-1* | 73 |
|  |  |  | *CDK2-2* | 89 |
| Related to Fig. S4f | FT282 hTERT *p53*<sup>R175H</sup> | WT | *PKMYT1-4* | 90 |
|  |  |  | *PKMYT1-6* | 88 |
|  |  | *CCNE1* (C3) | *PKMYT1-4* | 89 |
|  |  |  | *PKMYT1-6* | 84 |
| Related to Fig. S5j-k | FT282 hTERT *p53*<sup>R175H</sup> | *CCNE1* (C3) | *LMNA1-1* | 81 |

**Supplementary Table 4.**
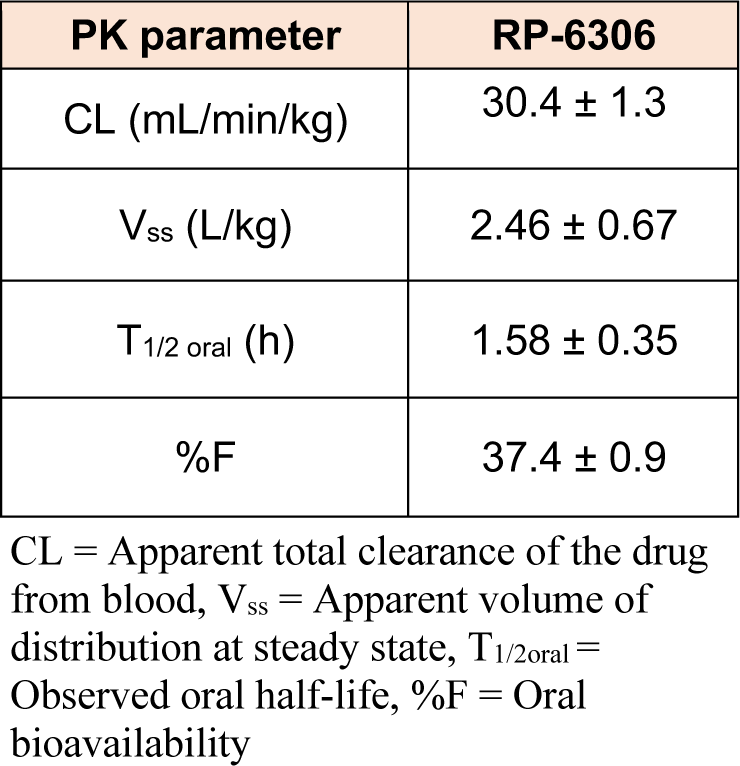
Pharmacokinetic parameters of RP-6306 in mice

**Supplementary Table 5.**
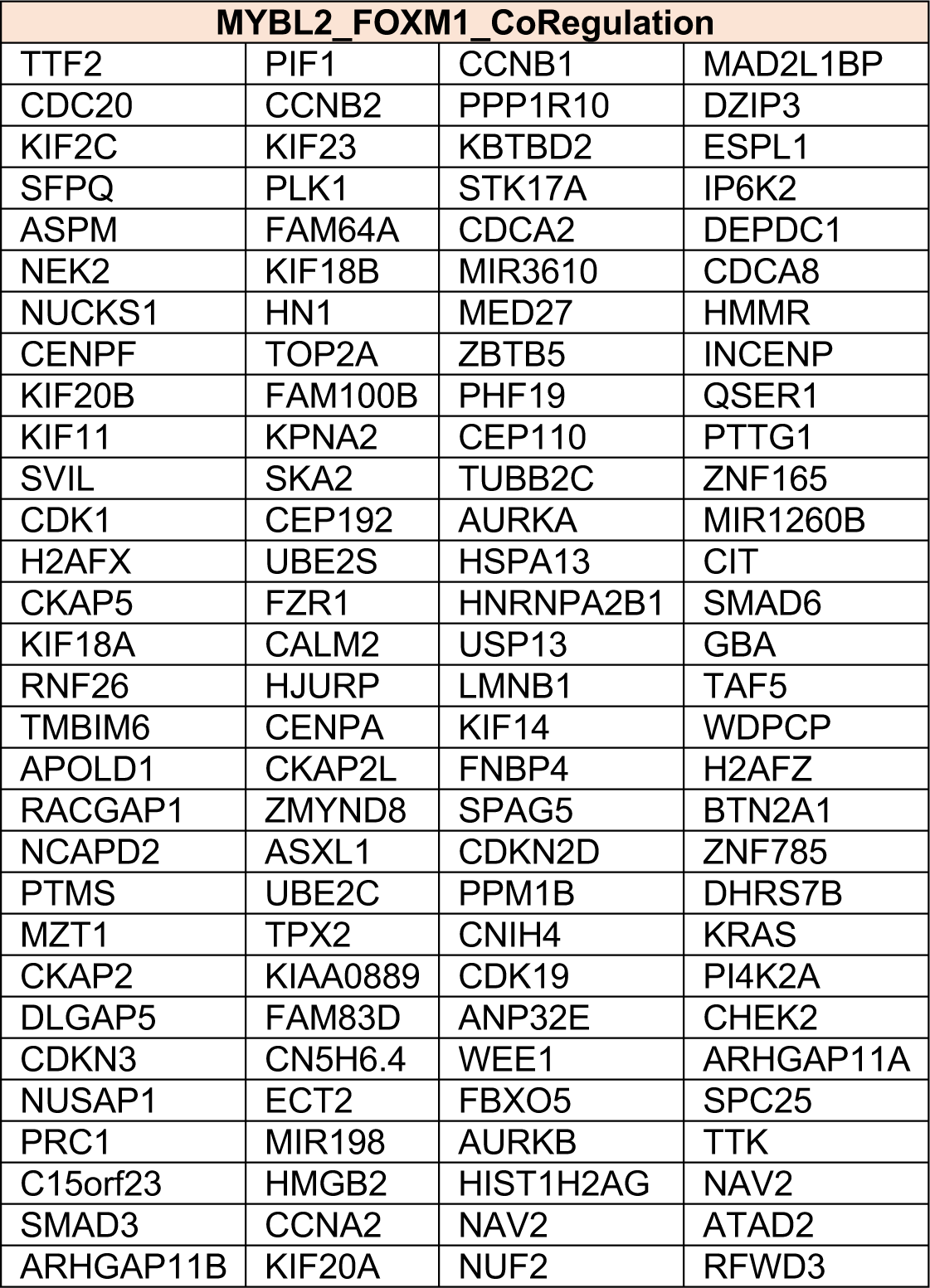
MYBL2-FOXM1 co-regulated genes used for GSEA analysis

| **MYBL2_FOXM1_CoRegulation** |  |  |  |
| --- | --- | --- | --- |
| TTF2 | PIF1 | CCNB1 | MAD2L1BP |
| CDC20 | CCNB2 | PPP1R10 | DZIP3 |
| KIF2C | KIF23 | KBTBD2 | ESPL1 |
| SFPQ | PLK1 | STK17A | IP6K2 |
| ASPM | FAM64A | CDCA2 | DEPDC1 |
| NEK2 | KIF18B | MIR3610 | CDCA8 |
| NUCKS1 | HN1 | MED27 | HMMR |
| CENPF | TOP2A | ZBTB5 | INCENP |
| KIF20B | FAM100B | PHF19 | QSER1 |
| KIF11 | KPNA2 | CEP110 | PTTG1 |
| SVIL | SKA2 | TUBB2C | ZNF165 |
| CDK1 | CEP192 | AURKA | MIR1260B |
| H2AFX | UBE2S | HSPA13 | CIT |
| CKAP5 | FZR1 | HNRNPA2B1 | SMAD6 |
| KIF18A | CALM2 | USP13 | GBA |
| RNF26 | HJURP | LMNB1 | TAF5 |
| TMBIM6 | CENPA | KIF14 | WDPCP |
| APOLD1 | CKAP2L | FNBP4 | H2AFZ |
| RACGAP1 | ZMYND8 | SPAG5 | BTN2A1 |
| NCAPD2 | ASXL1 | CDKN2D | ZNF785 |
| PTMS | UBE2C | PPM1B | DHRS7B |
| MZT1 | TPX2 | CNIH4 | KRAS |
| CKAP2 | KIAA0889 | CDK19 | PI4K2A |
| DLGAP5 | FAM83D | ANP32E | CHEK2 |
| CDKN3 | CN5H6.4 | WEE1 | ARHGAP11A |
| NUSAP1 | ECT2 | FBXO5 | SPC25 |
| PRC1 | MIR198 | AURKB | TTK |
| C15orf23 | HMGB2 | HIST1H2AG | NAV2 |
| SMAD3 | CCNA2 | NAV2 | ATAD2 |
| ARHGAP11B | KIF20A | NUF2 | RFWD3 |

**Supplementary Table 6.**
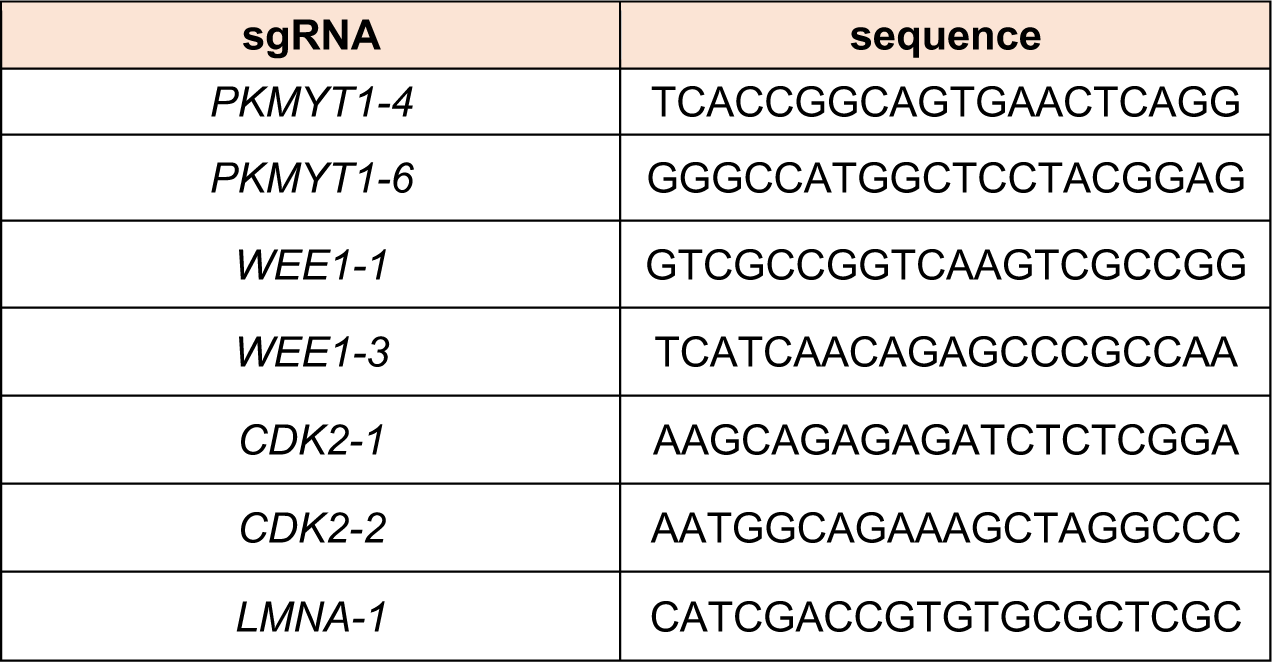
sgRNA guide sequences

